# Comparative Analysis of Salinity Response Transcriptomes in Salt-Tolerant Pokkali and Susceptible IR29 Rice

**DOI:** 10.1101/2023.08.20.551368

**Authors:** Matthew Geniza, Samuel E. Fox, Abigail Sage, Mitra Ansariola, Molly Megraw, Pankaj Jaiswal

## Abstract

Rice is a major cereal crop responsible for feeding the world’s population. To improve grain yield and quality, meet growing demand, and face the challenges posed by abiotic and biotic stress, it is imperative to explore genetic diversity in rice for candidate genes and loci that may contribute to stress tolerance. High salinity abiotic stress in the rice growth environment affects growth, yield, and quality. Therefore, we conducted a salt stress-responsive RNA-Seq-based transcriptome study of two rice (*Oryza sativa*) varieties, the salt-tolerant Pokkali and the salt-sensitive breeding line IR29. To identify early and late salinity response genes, we collected samples from the treated and untreated plants in this study at 1, 2, 5, 10, and 24 hours after treatment with 300 mM NaCl solution. We identified 7,209 and 6,595 salt-induced differentially expressed transcripts from Pokkali and IR29, respectively, over all time points. We identified ∼190,000 single nucleotide polymorphism (SNP) sites and ∼40,000 simple sequence repeat (SSR) sites, allowing analysis of their consequences on genetic diversity, transcript structure, gene function, and differential expression. We identified and validated the polymorphic SSRs in the differentially expressed salt-responsive genes *Respiratory Burst Oxidase Homolog B (RBOHB)* and *Rice Salt Sensitive 1 (RSS1)* that underly nearby salt tolerance QTLs. This study provides insight into transcriptional programming during salt stress, evidence for improving *Oryza* genome annotations, and reveals SNP and SSR sites associated with differential gene expression and potential gene function.

## Introduction

Rice is one of the most important cereal crops, and according to the International Rice Research Institute (IRRI), rice feeds more than 3.5 billion people—nearly 50% of the world’s population [1]. By 2050, the world population is projected to reach 9 billion, which means that rice production must increase to meet global demand. Therefore, developing and improving rice varieties that provide high grain quality and yield while also exhibiting resistance and tolerance to abiotic and biotic stresses is essential. An integrated genetics and genomics approach is becoming increasingly necessary to develop robust strategies to help breeding programs identify molecular markers closely associated with gene function and desirable phenotypes, such as salt tolerance.

Excess exposure to salt in coastal areas or due to saline soil/groundwater in the rice paddy growing environment adversely affects grain yield and quality. Salt-affected soil is prevalent in many of Asia’s regions, especially irrigated land [2]. The initial response to high salt conditions is hyperosmotic stress followed by rapid water loss in leaves [3]. An imbalance of Na^+^ and K^+^ ions in plants limits their ability to grow and develop by affecting physiological and cellular processes [4]. For example, salt stress negatively impacts chlorophyll biosynthesis, essential for photosynthesis [5] and the physiological processes of stomatal conductance and transpiration [6]. Reactive oxygen species are induced and can lead to oxidative damage to cells and disrupt cellular processes [7, 8]. The stage most sensitive to salt exposure are the early seedling and the reproductive stages of growth and development; however, different varieties may exhibit varied responses [9]. Only a handful of lines, such as Pokkali, that display tolerance to salt stress [9] provide potential gene targets that may confer salt tolerance in breeding lines.

We conducted a time course transcriptome study on rice seedlings treated with salt stress to explore and analyze differential gene expression, functional annotations, and potential consequences of genetic polymorphism on gene structure and function. The salt stress experiment was conducted on salt-tolerant landrace Pokkali and salt-sensitive breeding line IR29 varieties. We collected the seedling shoot samples from three biological replicates at 1, 2, 5, 10, and 24-hour time points from control plants (untreated) and plants exposed to 300 mM NaCl solution. Using computational tools and publicly available databases, we performed functional annotation of our transcriptome assemblies from the two rice lines. We observed time course-dependent transcriptome differences in response to salt stress in Pokkali and IR29 rice varieties. We identified previously known salt-responsive genes and a set of novel genes that were differentially expressed in the two rice lines. Using sequenced transcriptomes allowed us to identify SNPs (including indels) and SSRs as potential genetic markers colocalized in the genes’ transcribed regions. Therefore, this study provides important new resources for investigating the consequences of sequence polymorphism on the function, structure, and expression of the targeted gene sets for developing salt-tolerant breeding lines.

## Materials and Methods

### Plant material, growth conditions, and salinity treatment

Seeds of the *Oryza sativa* Indica group varieties salt sensitive/susceptible IR29 (USDA_GRIN: PI 393986) and salt tolerant Pokkali (USDA GRIN: GSOR 312020) were sown and watered thoroughly. The germinated seedlings continued growing under a 14 hr long photoperiod and room temperatures at 28°C during the day and 24°C at night. Two week old plants were treated with 300 mM NaCl solution, and the corresponding control plants were watered normally. One hour following salt treatment, tissue sampling began. We collected Three biological replicates of aboveground tissue (seedling shoot) of salt-exposed and untreated control plants were collected at 1, 2, 5, 10, and 24 hours post-salt exposure time points. Collected samples were immediately frozen in liquid nitrogen and stored at −80°C. Unlike the typical 0 hr time point in most time course studies, samples from untreated plants at each time point were collected as controls to minimize the effects of experimental conditions, physiology, and developmental processes occurring during the day of the experiment. The original seeds procured from the USDA GRIN germplasm repository were multiplied using a single seed descent prior to setting up the experiment.

### Sample preparation for sequencing

Total RNA from frozen seedling shoot samples was extracted using RNA Plant reagent (Invitrogen Inc., USA) and RNeasy kits (Qiagen Inc., USA) and treated with RNase-free DNase (Life Technologies Inc., USA). The purified mRNA fraction concentration and quality were determined using an ND-1000 spectrophotometer (Thermo Fisher Scientific Inc., USA) and Bioanalyzer 2100 (Agilent Technologies Inc., USA). Samples were prepared using the TruSeq^TM^ RNA Sample Preparation Kits (v2), and strand-specific sequencing was carried out on the Illumina HiSeq 2000 instrument (Illumina Inc., USA) at the Center for Genomic Research and Biocomputing, Oregon State University.

### De novo transcriptome assembly and annotation

FASTQ file generation from Illumina sequences was performed by CASAVA software v1.8.2 (Illumina Inc.). Illumina sequences were further filtered for low quality at a score of 20 using Sickle v1.33 [10]. It resulted in 907,218,339 and 911,926,822 51 bp reads to be used in the assembly process of IR29 and Pokkali, respectively. The transcripts were assembled using Velvet [11] (v1.2.10), which uses De Bruijn graphs to assemble short reads. An assembly of 31 and 37 k-mer lengths was performed separately for IR29 and Pokkali reads. Oases [12] (v0.2.08) produced transcript isoforms using sequence reads and pairing information and merging the assemblies produced by Velvet into a single consensus assembly. FullLengtherNext [13] was used to determine assembled transcripts with protein-coding capability, and tRNAscan-SE [14] (v1.3) was used to identify tRNA genes in our assemblies.

BLASTn [15] (E-value ≤ 1e^-10^) searches were performed with our assembled transcripts as a query against the reference genome and annotated cDNA sequence databases of *Oryza sativa japonica* (Japonica: Gramene IRGSPv1.0.21 and MSUv7.0) [16, 17], *O. sativa indica* (Gramene ASM465v1.21), *O. nivara* (Gramene: AWHD00000000.22) [18, 19], and the *O. sativa AUS* rice variety Kasalath [20] available from the Gramene database [18, 21].

The assembled transcripts were translated into the longest predicted open reading frame (ORF) using the ORFPredictor [22]. We used a combined approach based on functional motif analysis, sequence homology, Blast2GO [23], and BLASTx search (E-value ≤ 1e^-2^) against the NCBI GenBank nonredundant protein database [24, 25] to annotate the ORFs. The Yekutieli correction for false discovery rate calculation with significant cutoffs at a P value of 0.05 and a minimum of 5 mapping entries per GO term were applied for GO enrichment analysis using the AgriGO Analysis Toolkit [26].

### Gene expression analysis

CASHX v2.3 [27] aligned Illumina reads from IR29 and Pokkali to the reference Japonica gene models, and normalized counts were determined (Supplementary Table 1). The EdgeR package v3.3.2 [28] was used to determine significant differential gene expression. Groups were determined by variety (IR29 or Pokkali), treatment (Control or Salt), and time point (1 hr, 2 hr, 5 hr, 10 hr, or 24 hr post-treatment). Differentially expressed transcripts were filtered for significance P value cutoff/False Discovery Rate Corrected P value of 0.05. Furthermore, differentially expressed transcripts were filtered by log (2)-fold cutoffs. Multidimensional scaling (MDS) was used to assess and visualize a cross-sample comparison. MDS shows clustering based on count values for all transcripts among all replicates (Supplementary Figure S1).

### Transcription factor gene analysis

All differentially expressed genes were cross-referenced against the rice transcription factor database hosted by Grassius [29] to explore the expression of transcription factors in our dataset. The subset of differentially expressed transcription factors was clustered using Pearson correlation with an r-value of 0.7 in BioLayout Express3D [30]. To find the transcription factor-binding sites (TFBS) and co-expressed transcription factors potentially regulating the salt-responsive genes, the TFBS scanner suite [31] was used to calculate the likelihood of transcription factors whose binding domain is represented as a Positional Weight Matrix (PWM) to bind to a promoter region of a gene of interest. The binding site scan was carried out on the 1 kb upstream region of the genes of interest (Supplementary Figure S2), and log-likelihood scores for each location were computed. A stringent false positive rate of 0.0001 (0.01%) was used to report the putative TFBSs for each PWM.

### Genetic marker development

SSRs were identified in the IR29 and Pokkali assemblies by using the stand-alone version of the Simple Sequence Repeat Identification Tool (SSRIT) [32]. To identify SNPs, an alignment database of the *Oryza sativa japonica* genome was created by Spliced Transcripts Alignment to a Reference (STAR v2.4.1d) [33]. Illumina FASTQ reads were aligned to the database using STAR. The generated alignments were sorted using samtools [34]. Sorted alignments were run using VarScan (v2.3.9) (--min-coverage 2 --min-var-freq 0.8 --p value 0.005) [35], and output files were reformatted to VCF output and run through the Ensembl Plants API Effect Predictor tool [36] to obtain potential consequences of the identified SNPs (Variant Effect Prediction analysis using resources available at Gramene database).

Primers for rice genes *RBOHB* (Forward: TCAAGCTGTCCAAACACCGT; reverse: GCGACCACAAAAAGCTGGAG) and *RSS1* (Forward: ACTCCTGGAGCCTGGAATGA; reverse: TTGCTTCCGCTACTTGGGTT) were developed using Primer3Plus [37] and evaluated for target specificity using NCBI Primer-BLAST [38]. The gene fragments were amplified from the genomic DNA from various rice lines, including the reference *O. sativa japonica* cv Nipponbare. PCR was carried out for 30 cycles. Each amplification cycle consisted of denaturation at 94°C for 15 seconds, annealing at 65°C for a minute, extension at 72°C for 30 seconds, and cooling at 4°C for 15 seconds. The amplified products from the respective rice lines were purified and sequenced on an Applied Biosciences 3730 gel capillary sequencer at the CGRB. The 3000 rice genomes data underlying the amplified regions of *RBOHB* and *RSS1* genes was downloaded from the Rice SNP-seek database [39] and the Croppedia online tool (Keygene Inc). Alignments of the sequenced regions of the *RBOHB* and *RSS1* genes were generated using the multiple sequence alignment software MUSCLE [40, 41]. The mRNA secondary structures were predicted by the RNAfold software accessed online from the ViennaRNA Web Services [42, 43].

### Data availability

Raw FASTQ files from the RNA-Seq experiment were submitted to EMBL-EBI ArrayExpress. They are accessible under the accession E-MTAB-1913 for IR29 and E-MTAB-1914 for Pokkali. Primer sequences used to amplify SSR regions of *RBOHB* (ProbeDB_acc: Pr032825000) and *RSS1* (ProbeDB_acc: Pr032825001) are available at the NCBI Probe Database. Nucleotide sequences from amplified fragments of *RBOHB* and *RSS1* from multiple rice lines, as mentioned in the plant material, are available from EMBL-EBI ArrayExpress accessions E-MTAB-5560 (*RBOHB*) and E-MTAB-5583 (*RSS1*).

## Results

### Transcriptome Sequencing

In all, 60 cDNA libraries were generated and sequenced from shoot tissues of 3 biological replicates of IR29 and Pokkali at 1, 2, 5, 10, and 24 hours post-salt treatment and their respective time point control from the untreated plants. The 51 base pair single-end sequencing of the cDNA libraries on the Illumina HiSeq 2000 resulted in 161 Gb and 163 Gb of nucleotide sequences for IR29 and Pokkali, respectively.

### Overview of differential expression in response to salt treatment

High-quality filtered RNA-Seq reads from IR29 and Pokkali were used for the transcriptome analyses. To examine salt-induced gene expression, the 51 bp reads from the control untreated and salt-treated samples from each of the five-time points generated by the Illumina HiSeq 2000 were aligned to the *O. sativa japonica* genome (IRGSP-1.0.26; japonica rice genome). The Japonica rice genome was chosen for alignment for its high-quality genome assembly and annotation. Across all time points, 4336 genes were upregulated, and 2259 genes were downregulated in the salt-susceptible IR29 compared to the salt-tolerant variety Pokkali, where 3821 upregulated and 3388 downregulated genes were observed. The two varieties shared 2501 differentially expressed genes, of which 140 were differentially expressed across all time points (Figure 1A).

**Figure 1.**
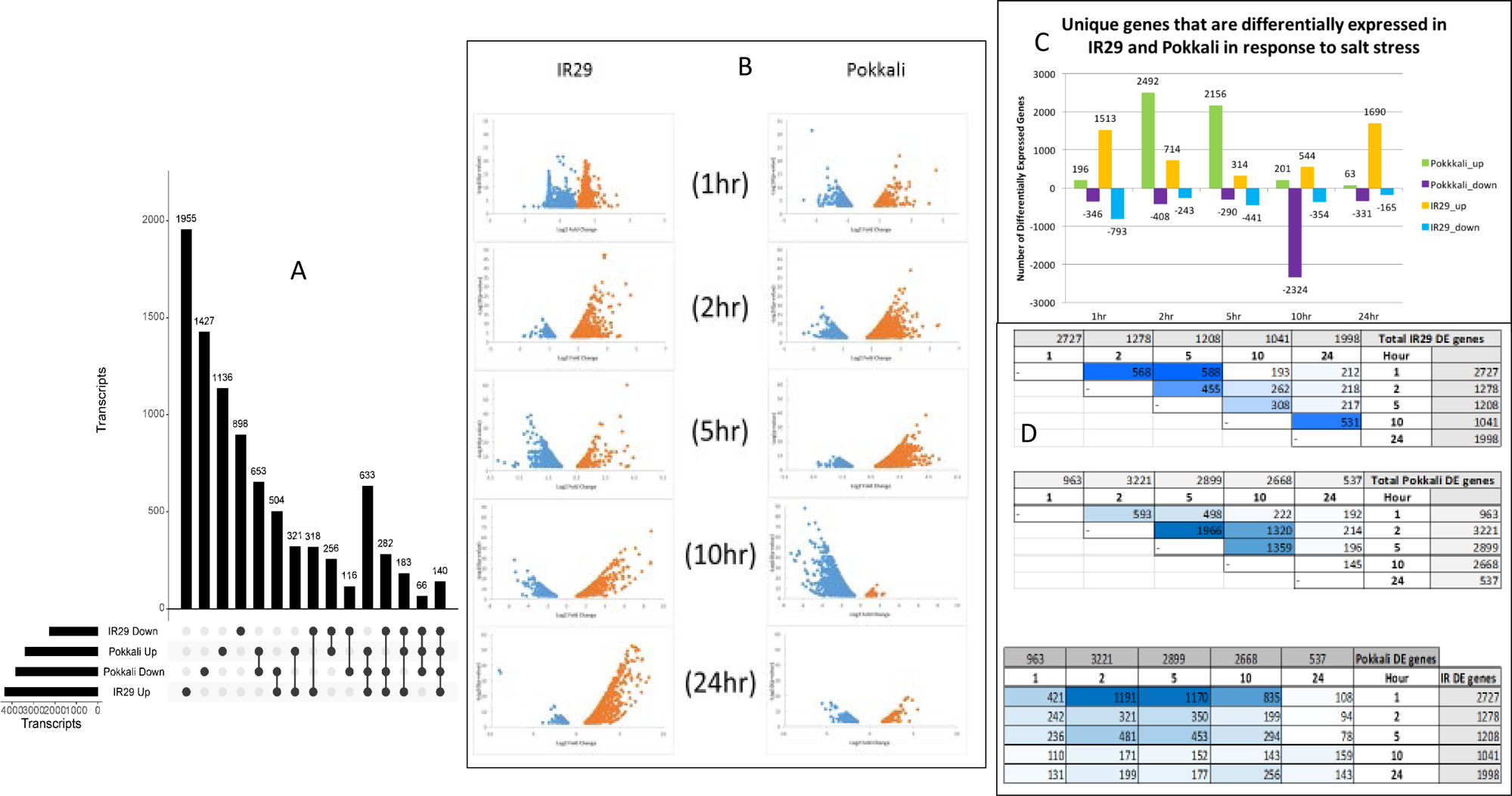
A. Analyses of differentially expressed transcripts over 5 time points. Upset plot show the distribution and commonalities of Japonica homolog counts with respect to their response post salt exposure in IR29 and Pokkali. B. Analyses of differentially expressed transcripts over 5 time points. A scatter plot of light upregulated (orange-colored) and downregulated (blue-colored) transcripts from IR29 and Pokkali. Each point represents a single transcript. C. The bar graph shows the number of unique transcripts that are differentially expressed at each time point in IR29 and Pokkali. D. Tables reveal the number of transcripts in common between time points in each variety (IR29 – top, Pokkali – middle) and the number of transcripts in common between time points between each variety (bottom).

### Hour-by-hour gene expression in response to salt treatment

Transcriptome data from 5 separate time points per variety per treatment enabled us to capture the variety- and time-specific plant response to salt stress. Unlike classical time series experiments using a zero timepoint, we performed the gene expression analysis by comparing the untreated sample from the same time point to avoid any biases resulting from response to diurnal regulation. IR29 had a total of 2727, 1278, 1208, 1041, and 1998 differentially expressed transcripts at 1, 2, 5, 10, and 24 hours, respectively, compared to Pokkali’s 963, 3221, 2899, 2668, and 537 differentially expressed genes at the corresponding time points (Figure 1B and 1C, Supplementary Figure S3). In IR29, we observed an immediate response to salt treatment, as 2,727 genes were determined to be differentially expressed (up- or downregulated) at the 1-hour time point (Figure 1D - Top). The number of genes was reduced by half in hours 2 and 5, and nearly 50% were regulated in hour 1. At 24 hours, the overall number of differentially expressed genes represented most of the expression patterns observed at hour 1; however, only 1/10 of the same genes were present at hour 1. Approximately 50% of the genes at hour 10 remained differentially expressed at 24 hours.

Compared to IR29, salt-tolerant Pokkali carries fewer differentially expressed genes at hour 1 and an increasing number at hours 2, 5, and 10. Approximately 50% of the genes exhibiting differential expression during hours 2-10 were the same at each time point, indicating a consistent genetic response in Pokkali during these hours. At 24 hours, the number of differentially expressed genes decreased to a level similar to that at hour 1. Approximately 35% of the genes that were up- or downregulated at 24 hours were also expressed at hour 1 (Figure 1D - Middle).

When we compared the gene expression profiles at each time point between the two varieties, we observed that the genetic response to salt stress was quite different – very few of the same genes were similarly expressed between the two varieties at each time point. An exception to this is the most likely initial response to salt stress. Greater than 1/3 of the genes expressed in IR29 at hour 1 were expressed in Pokkali at hours 2 and 5 (Figure 1D - Bottom). Examining the differentially expressed genes, Pokkali most likely cycled and acclimated, while IR29 expressed 4 times as many genes at 24 hours post-salt exposure. A multidimensional scaling (MDS) analysis using the read counts of aligned transcripts of IR29 or Pokkali samples corroborates expression profiling, and samples from time points that display similar expression patterns are clustered closer together (Supplementary Figure S1A-B). When comparing all samples against each other, we observed the formation of 2 distinct clusters that represented the IR29 and Pokkali varieties (Supplementary Figure S1C). While we analyzed the transcriptome data at the genome scale, we also looked at the expression of individual genes known for their salt responsiveness (Supplementary Figure S2). Among these genes, *OSBZ8* (OS01G0658900), a bZIP class *ABSCISIC ACID RESPONSE ELEMENT-BINDING FACTOR,* is highly expressed in salt-tolerant cultivars as compared to salt-sensitive cultivars [44]. Our analysis revealed that although *OSBZ*8 was upregulated in both lines, Pokkali showed a 4-fold change at 24 hours compared to only a 2-fold change in IR29 at 24 and 10 hours. The AP2-EREBF/DREB transcription factor family gene (OS04G0549700) is known to be induced by cold stress [45]. Since both cold stress and salt stress share a response by stimulating the collection of osmolytes and antioxidants to interact with stress-induced free radicals [46], we expected such genes to be differentially expressed in our study.

Similarly, plasma membrane intrinsic protein family members (*PIP*, OS04G0233400, OS09G0541000), Nod26-like intrinsic proteins (*NIP*, OS02G0232900, OS06G0228200), and small and basic intrinsic proteins (*SIP*, OS07G0209100) were upregulated in both IR29 and Pokkali in response to salt stress. These proteins are subfamilies of plant aquaporins, which are extensively involved in regulating physiological features during plant growth and development and salt tolerance [47]. PIPs were upregulated 1.6-fold at 1 hour in IR29, whereas in Pokkali, they were upregulated by at least 2.5-fold at 2 and 5 hours and downregulated at 10 hours. *NIP* follows a similar trend to *PIPs*. *SIP* was upregulated at 10 and 24 hours in IR29 and at 24 hours in Pokkali.

Specific to salt tolerance, we observed differential expression in *Rice Salt Sensitive 1* (*RSS1*, OS02G0606700). *RSS1* is involved in the cell cycle, and its expression confers salt tolerance [48]. In Pokkali, *RSS1* was upregulated at 1, 2, 5, and 24 hours and downregulated at 10 hours. In IR29, *RSS1* was downregulated at both 5 and 10 hours. The expression of the gene encoding the late embryogenesis abundant (*LEA*) protein is associated with increased structural stability and improved stress tolerance in transgenic plants [49]. *LEA* protein-coding genes are upregulated late in IR29 at 24 hours as compared to early expression at 1 hour in Pokkali. Overexpression of the major Na^+^ transporter *HKT1* is known to confer salt tolerance in wheat [4]. Its rice homolog *OsHKT1* (OS06G0701700) was found to be upregulated in Pokkali at 2 hours but not significantly expressed in IR29. The known rice salt-responsive gene *NUC1* [50] (OS08G0192900) shows a similar profile in both lines, except it is downregulated in IR29. Its overall expression is much higher in Pokkali at 2, 5, and 10 hours.

### Expression profiles of transcription factor gene family members

Using publicly available transcription factor databases [51], we identified 271 and 285 transcription factor genes in IR29 and Pokkali that were differentially expressed at one or more time points. A total of 424 different transcription factors were determined to be differentially expressed in this study (Supplementary Table 2). Of these 424 transcription factors, only two, OS01G0313300, an ethylene response factor (*ERF*) described above, and OS03G0335200, a *WRKY* [52], were differentially expressed at all time points (Supplementary Figure S2). The 424 transcription factors represented 36 Panther gene families. AP2-EREBP, MYB, WRKY, and NAC were the most represented transcription factor families. The AP2-EREBP/DREB family member as described above. MYB transcription factors are well studied for their broad involvement in regulating defense responses to biotic and abiotic stresses, hormone signaling, and many metabolic and developmental processes in plants [53–55]. By enhancing the transduction of stress response signals, WRKY and NAC proteins are known to regulate stomatal aperture [56, 57]. A coexpression analysis revealed 11 correlated clusters of differentially expressed transcription factors (Figure 2A). Most clusters were unique to rice variety and expression pattern (Pokkali Clusters 1, 5, 7, 11, and IR29 Clusters 2, 4, 8). Clusters 2, 6, 9, and 10 contained differentially expressed transcription factors in both rice varieties (Figure 2B). Of the 424 differentially expressed transcription factors in this study, 129 were found to be diurnally regulated by querying against the Diurnal database [58].

**Figure 2.**
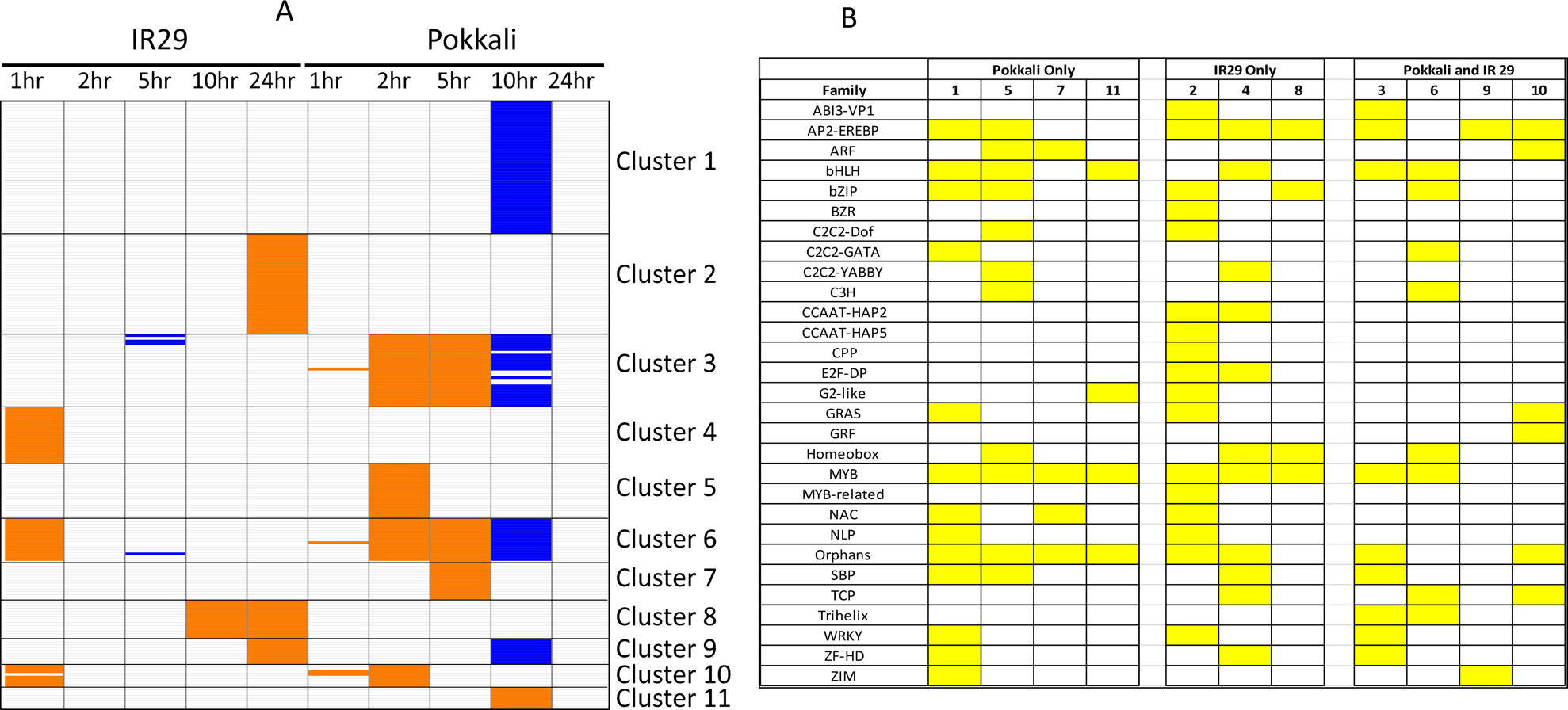
A. Matrix of transcription factors differentially expressed in response to salt exposure during the time course. The columns contain the variety and respective time point. Each cluster row width represents the number of transcripts for a given set of transcription factors. Orange color shows time-specific upregulated expression, and blue color shows time-specific downregulated expression. B. List of transcription factor families and presence of their transcripts in a given cluster.

Most transcription factors within each cluster shared a similar gene expression profile reported in the rice variety IR64 [59]. A comparative view of clusters reveals rice variety, transcription factor family, and time point-specific expression. In Pokkali-specific clusters 1, 5, 7, or 11, we observed differential expression of Auxin Response Factors (ARF), Jasmonate ZIM-Domains (ZIM), and C2C2-GATA transcription factors. The ZIM and C2C2-GATA transcription factors share the TIFY domain and are known to have important roles in abiotic stress [60]. Differentially expressed transcription factors exclusive to IR29 (clusters 2, 4, or 8) include abscisic acid insensitive (ABI) and brassinosteroid regulation (BZR) family members.

ABI transcription factors are known to be induced by salt stress, and their overexpression confers high sensitivity to salt stress [61]. The *OsBZR1* in cluster #2 was upregulated in the developing seedlings of the IR20 rice variety under drought stress (EMBL EBI ArrayExpress Experiment E-GEOD-41647). Based on the GO annotations, these clusters of transcription factors are involved in general biological processes; apoptosis, metabolic and developmental processes. Clusters 2 and 3 carry enrichment of transcription factors involved in response to a stimulus. Most of these genes showed regulation in IR29 at 24 hours compared to the salt-tolerant Pokkali at 2, 5, and 10 hours.

### De novo transcriptome assembly of IR29 and Pokkali

The de novo assemblies of the IR29 and Pokkali transcriptomes were created using [11] and Oases [12]. The assemblies of each cultivar were created in two steps: first, two separate assemblies were created from optimized 31 and 37 K-mer lengths; second, transcript isoforms that were assembled at both K-mer lengths were merged to represent the total number of unique transcripts in a consensus assembly of IR29 and Pokkali. It resulted in 109,407 and 106,134 transcripts ≥ 200 bp in length for IR29 and Pokkali, respectively.

The overall number of transcripts (the coverage of assembly) and average length and diversity of transcripts (the estimated number of discrete loci assembled) (Table 1) are the core statistics for the initial assembly quality assessment. The quality assessment software Full-LengtherNEXT [13] was used on the assemblies to classify transcripts as unigenes, determine if any transcripts had orthologs in databases, and suggest which unknown transcripts were protein-coding. Full-LengtherNEXT determined that 89% of the transcripts had orthologs for both assemblies. For the 10% of the assemblies that did not have orthologs, ∼33% of these transcripts were protein-coding. These assembled transcripts may include novel isoforms (Table 2).

**Table 1:**
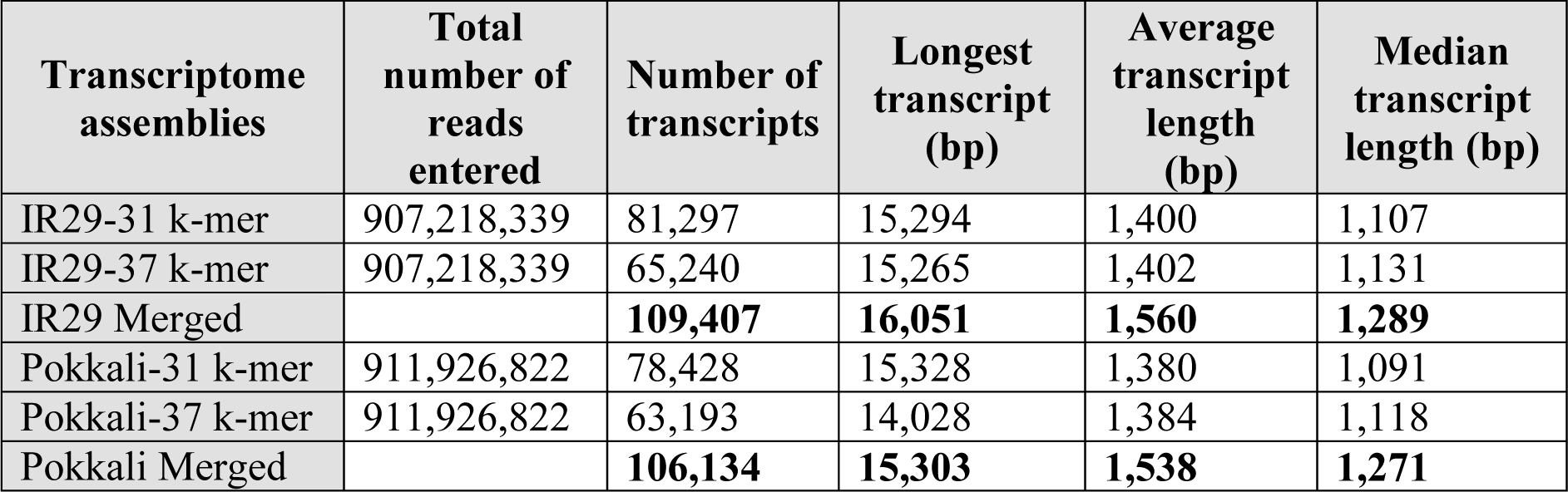
Transcriptome assembly statistics. Statistics for *Oryza sativa* Indica group varieties IR29 and Pokkali. Assemblies were generated by Velvet/Oases. The statistics describe the number of reads input to the assembler and the number of assembled transcripts and transcript lengths. The ability to merge assemblies via transcript isoforms into putative gene loci is a feature of Oases.

**Table 2:** Transcriptome assembly classification. Statistics for *Oryza sativa* Indica group varieties IR29 and Pokkali generated by Full-LengtherNEXT software classifying transcripts as unigenes, determining if any transcripts had orthologs in databases and suggesting which unknown transcripts were protein-coding.

|  |  | IR29 |  | Pokkali |  |
| --- | --- | --- | --- | --- | --- |
| Status |  | Transcripts | % | Transcripts | % |
| Complete | Sure | 34600 | 31.63 | 34631 | 32.63 |
|  | Putative | 10696 | 9.78 | 10241 | 9.65 |
| C-terminus | Sure | 21345 | 19.51 | 20843 | 19.64 |
|  | Putative | 3359 | 3.07 | 3044 | 2.87 |
| N-terminus | Sure | 14582 | 13.33 | 13446 | 12.67 |
|  | Putative | 4181 | 3.82 | 4002 | 3.77 |
| Internal |  | 8860 | 8.10 | 8282 | 7.80 |
| Misassembled |  | 463 | 0.42 | 478 | 0.45 |
| Putative Chimera |  | 0 | 0.00 | 0 | 0.00 |
| Coding | Sure | 1194 | 1.09 | 1268 | 1.19 |
|  | Putative | 2014 | 1.84 | 1921 | 1.81 |
| Putative ncRNA |  | 246 | 0.22 | 242 | 0.23 |
| Unknown |  | 7867 | 7.19 | 7736 | 7.29 |
| Total |  | 109407 | 100.00 | 106134 | 100.00 |
|  |  | IR29 |  | Pokkali |  |
| Unigene Report |  | Transcripts | % | Transcripts | % |
| with ortholog in the Database |  | 98086 | 89.58 | 94967 | 89.48 |
| Putative New Genes |  | 3208 | 2.93 | 3189 | 3.00 |
| ncRNA's |  | 246 | 0.22 | 242 | 0.23 |
| Unknown |  | 7867 | 7.18 | 7736 | 7.29 |
| Total |  | 109407 | 100.00 | 106134 | 100.00 |
| Without ortholog |  | 11321 | 10.12 | 11167 | 10.29 |
| Coding > 200 bp |  | 1194 | 10.55 | 1268 | 11.35 |
| Coding > 500 bp |  | 531 | 4.69 | 547 | 4.90 |
| Putative Coding > 200 bp |  | 2014 | 17.79 | 1921 | 17.20 |
| Putative Coding > 500 bp |  | 1047 | 9.25 | 1052 | 9.42 |

Furthermore, less than 0.5% of the total assemblies were potentially misassembled, and neither were considered chimeric. We observed that the de novo assembled transcripts were always longer than the reference Japonica transcript length in almost all size ranges, except for the 250 bp and 1000-1500 bp lengths (Supplementary Figure S4). We also identified 126 and 96 tRNA genes in the IR29 and Pokkali assemblies using tRNAscan-SE [14] (v1.3).

The Blast2GO [23] pipeline provided Gene Ontology (GO) annotations for IR29 and Pokkali transcripts, resulting in 75,236 (68%) and 72,686 (68%) unique transcripts that had at least one GO annotation for IR29 and Pokkali, respectively. Overall, 1,818 and 1,798 GO terms were assigned to IR29 and Pokkali, respectively, and 1,741 of these were shared between the two varieties. Seventy-seven GO terms were unique to IR29, and 57 were unique to Pokkali (Supplementary Figure S4; Supplementary Table 3). Of these transcription factors, binding activity, stress response, terpenoid biosynthesis, and cell cycle regulation were enriched in Pokkali. However, protein degradation, spliceosomal complex, chromosomal modification, and vesicular traffic terms enriched for IR29.

### Comparing transcriptome assemblies to other Oryza genomes

To annotate, characterize, and approximate the coverage of assembled transcripts and check for assembly quality, we compared the IR29 and Pokkali transcript assemblies with genomes and transcripts of the *Oryza* genus using BLAST (Table 3). Over 99% of assembled transcripts from both lines were mapped to the *O. sativa indica* and japonica reference genomes. At least 90% of our transcripts were mapped to annotated reference transcripts. We also compared our assembled transcripts to *O. nivara*, which shares a recent common ancestor of the AA genome with *O sativa* [62]. Approximately 99% and 94% of transcripts were mapped to the *O. nivara* genome and transcripts, respectively (Table 3). The recently sequenced genome and transcripts of the *O. sativa AUS* rice variety Kasalath [20] were also used in our comparisons.

**Table 3:**
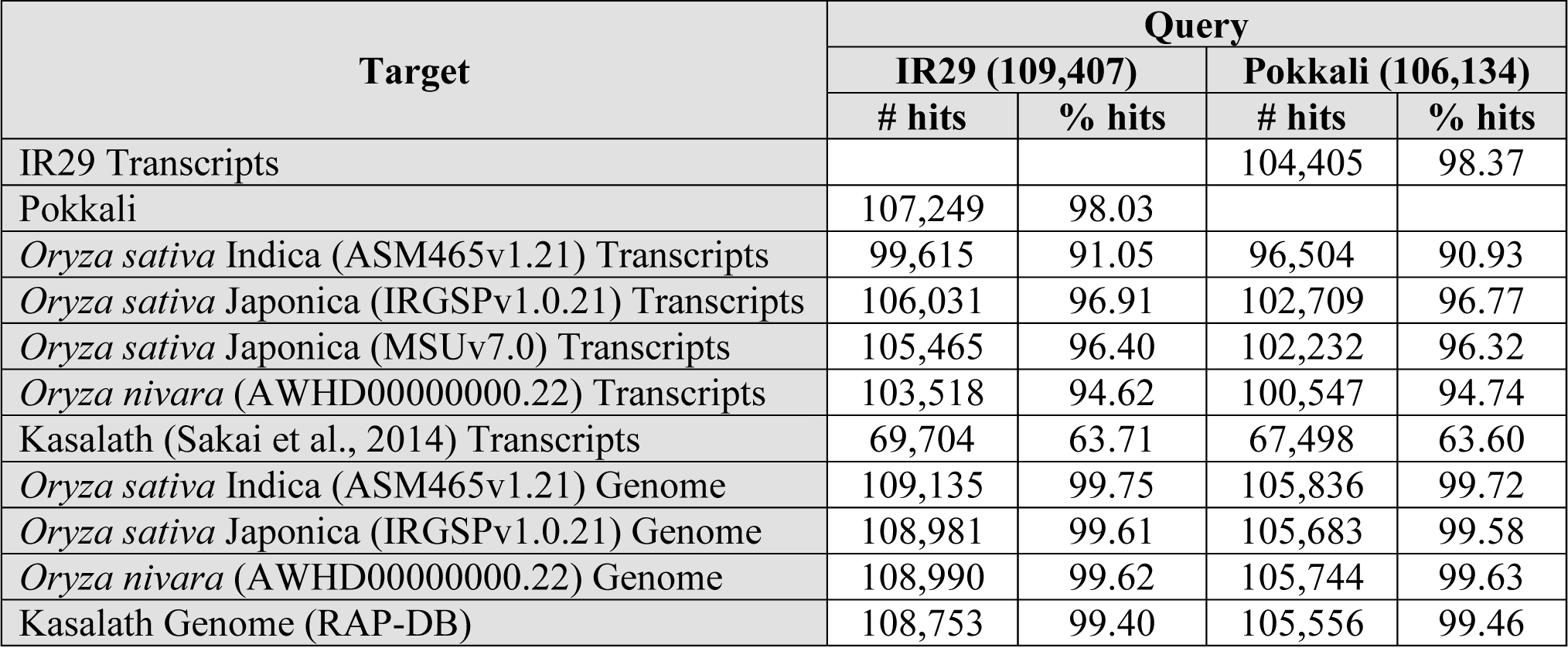
BLAST results. BLASTn (E-value 1e^-10^) nucleotide sequence comparisons of *Oryza sativa* Indica group varieties IR29 and Pokkali transcripts against gene models and genomes from other sequenced rice species suggested the coverage represented in the *Oryza sativa* Indica group varieties IR29 and Pokkali transcriptomes.

Although 99% of all transcripts mapped to the sequenced japonica reference genome, a small fraction of the 2,916 IR29 and 2,976 Pokkali transcripts did not map to annotated gene models. Using positional information from the genome (chromosome number and nucleotide start-stop obtained from the BioMart tool provided by the Gramene database [19]), we obtained sets of genes within the boundary of the assembled transcript. For example, in Pokkali, such assembled transcripts were found to hit regions near HOX-like protein coding the OS06G0140700 (*HOX2*) and OS10G0561800 (*HOX1*) genes. In IR29, transcripts hit regions of the OS05G0322900 (*WRKY*), OS02G0565600 (*HOX7*), and OS06G0140700 (*HOX2*) genes. HOX and WRKY gene family members affect stress response and plant development [63, 64]. Forty-eight of these genes in IR29 and 46 in Pokkali are differentially expressed in our analysis (Supplementary Table 4).

### Identification of genetic marker loci

To further advance our search for markers that may have associations with the salt response, we cross-referenced the genome position of all differentially expressed and the colocalized SSRs and SNPs sites, for example, on chromosome 1 (Figure 3).

**Figure 3.**
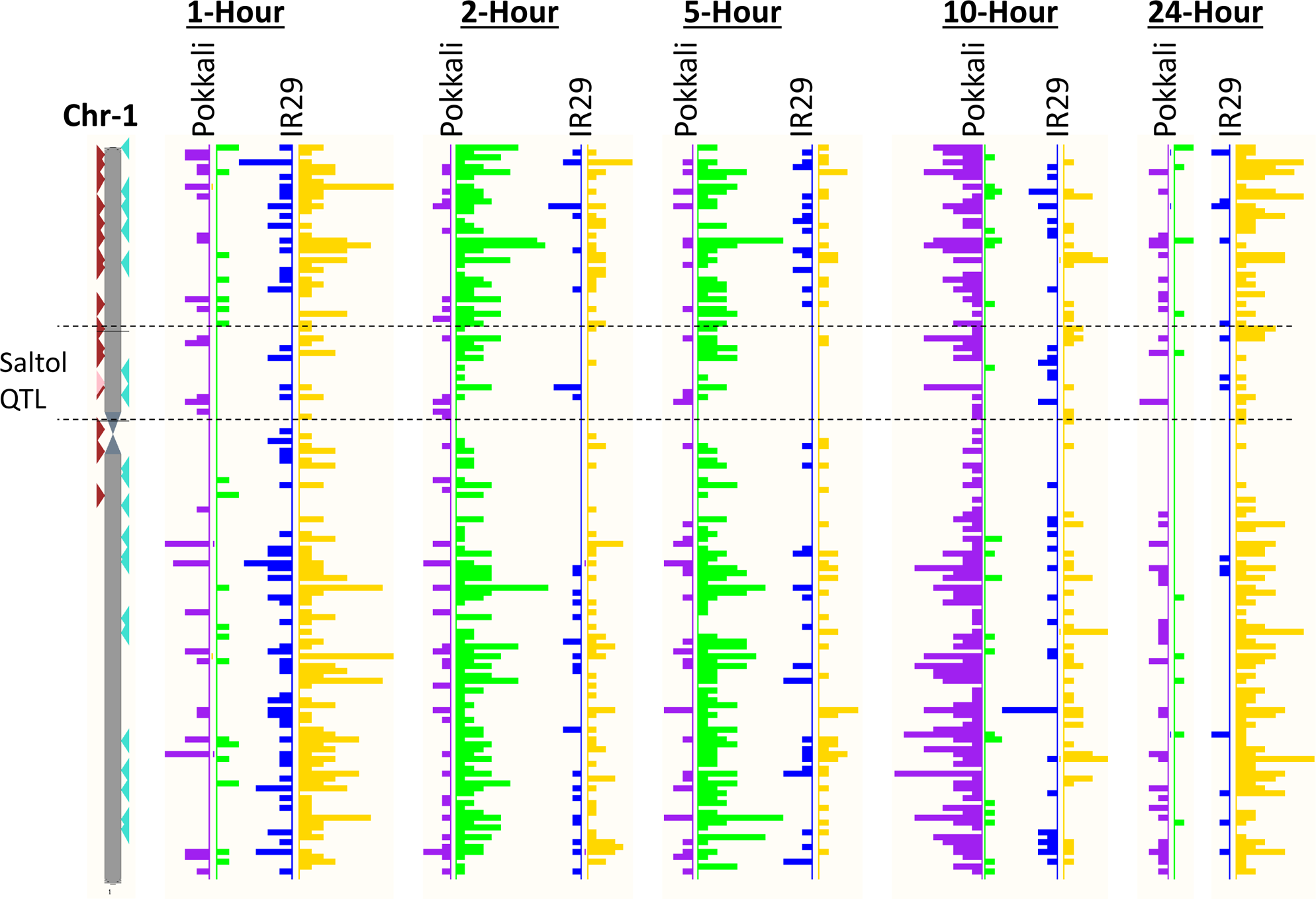
Fold change values of differentially expressed transcripts of IR29 and Pokkali mapped to chromosome 1 of the reference Japonica genome. Polymorphic SNPs and common polymorphic SSRs are also plotted to show markers that may be useful in selecting salt-stress traits. The saltol QTL region is marked between the dashed lines. The left side of each bar graph depicts upregulation.

### Simple sequence repeat (SSR) marker sites

Using the de novo assembled transcripts, we identified 42,693 SSRs in 29,919 unique transcripts of IR29 and 40,374 SSRs in 28,709 unique transcripts in Pokkali. These SSRs had dimer, trimer, tetramer, pentamer, and hexamer repeats (Supplementary Figure S6, Supplementary Table 5). We identified 4,656 homologous transcripts between IR29 and Pokkali that shared the same SSR motif. Seven hundred homologous transcripts overlapping 342 reference japonica gene sites bear the same SSR motif of polymorphic length. These 342 SSRs exhibit a uniform distribution throughout the Japonica genome, which makes them potential targets as genetic markers in rice breeding programs.

Many polymorphic SSR sites overlap the known abiotic stress tolerance QTLs and the differentially expressed genes identified in our dataset. For example, the differentially expressed gene *RBOHB* carrying a polymorphic dinucleotide (GA)n SSR in the 5’-UTR overlaps the QTLs associated with salt tolerance [65–68], potassium chlorate resistance (Gramene QTL ID#AQF091, #CQI1), sodium uptake (#CQI2), sodium to potassium ratio (CQI3), root penetration (CQAW4), and lodging incidence (#AQDZ002, #AQDZ013). Similarly, the gene *RSS1* carrying a trinucleotide (GAT)n SSR in the 3’-UTR is known to confer salt tolerance; it overlaps QTLs associated with osmotic maintenance (#AQDX004), root penetration (#AQC005, #DQF1), and drought tolerance traits [69–73]. We validated the polymorphic SSRs found in the UTR regions of *RBOHB* and *RSS1* by sequencing the amplified genomic regions from IR29, Pokkali, and a handful of indica and japonica rice accessions. The sequences obtained were aligned to the reference japonica genome and the wild *Oryza* and ancestor genomes to explore the diversification of the SSR sites. Observations from this analysis are discussed later.

### Single nucleotide polymorphism (SNP) marker sites

We identified more than 282,618 potential SNP sites in IR29 and Pokkali by aligning reads to the japonica reference genome with an average of one SNP per 3,179 bp of the assembled genome (Supplementary Figure S7). Of the 117,762 sites common to both varieties, 257 sites bear different alleles for IR29 and Pokkali (Figures 3 and 4, Supplementary Table 6). IR29 contained more nucleotide transitions and transversions (Supplementary Table 7). Using the Variant Effect Prediction (VEP) tool [36], we also predicted the potential consequences of these variants on the regulatory region, structure, splicing, and function of the genes (Table 4).

**Figure 4.**
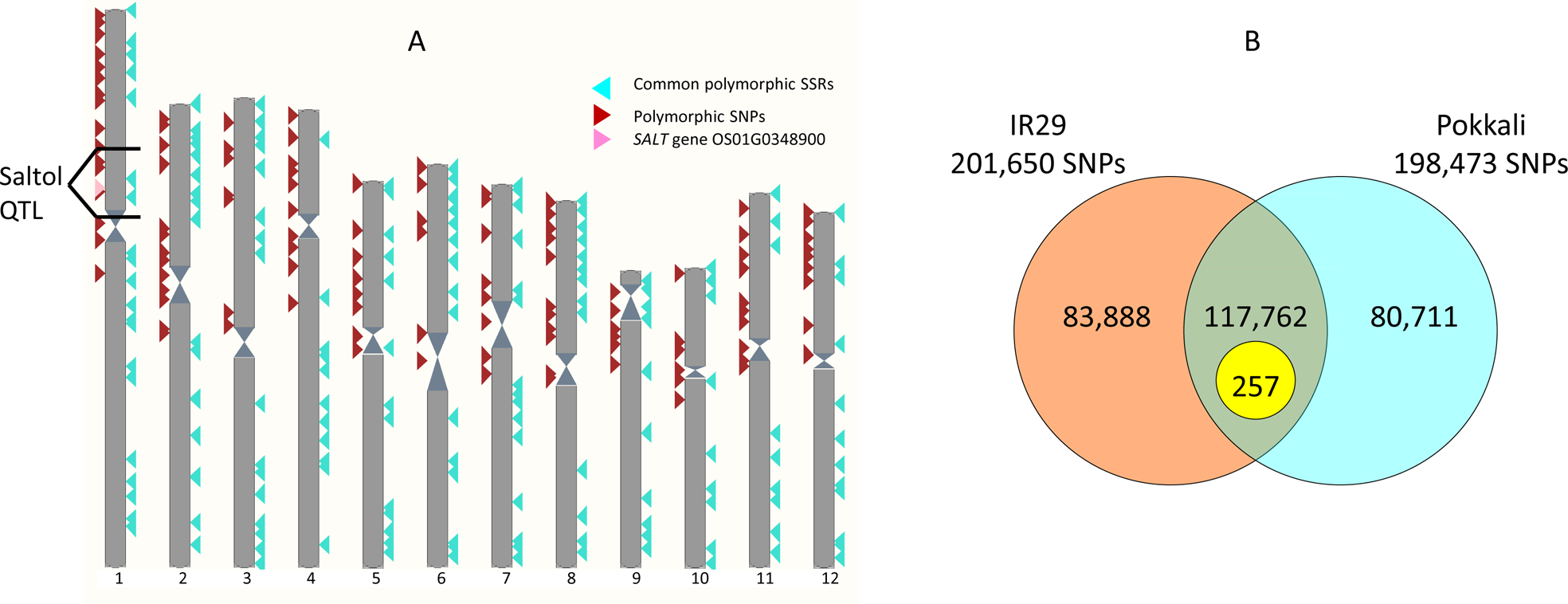
A. Genetic marker discovery. Mapping of the 257 polymorphic SNPs (left side, brown triangle) and 700 common polymorphic SSRs (right side, aqua triangle) in the IR29 and Pokkali transcriptomes on the karyotype view of the reference Japonica genome hosted by Ensembl Plants. The region of the Saltol QTL and *SALT* stress-induced gene (OS01G0348900) are also highlighted on the karyotype view. B. Number of SNPs identified in IR29 and Pokkali after alignment against the reference Japonica genome. A total of 257 common SNP sites contained polymorphic alleles in IR29 and Pokkali compared to the reference Japonica allele.

**Table 4;.** Prediction of SNP variant consequences. SNP variant consequence prediction based on IR29 and Pokkali SNPs was identified by aligning the sequenced reads from IR29 and Pokkali to the reference Japonica genome available from the Ensembl Plants database. Listed variant effect types are based on the categories adopted by the Ensembl Plants database.

| Predicted variant effect | Number of SNP sites with consequences |  | Unique |  |
| --- | --- | --- | --- | --- |
|  | IR29 | Pokkali | IR29 | Pokkali |
| splice donor variant | 3729 | 3810 | 549 | 630 |
| splice acceptor variant | 2577 | 2587 | 391 | 401 |
| stop gained | 335 | 360 | 145 | 170 |
| frameshift variant | 199 | 221 | 196 | 218 |
| stop lost | 174 | 180 | 65 | 71 |
| start lost | 85 | 73 | 40 | 28 |
| inframe insertion | 23 | 26 | 23 | 26 |
| inframe deletion | 160 | 161 | 160 | 161 |
| protein altering variant | 3 | 2 | 3 | 2 |
| missense variant | 13070 | 13172 | 3219 | 3321 |
| splice region variant | 5125 | 5208 | 995 | 1078 |
| incomplete terminal codon variant | 3 | 3 | 1 | 1 |
| synonymous variant | 12823 | 12823 | 3123 | 3123 |
| stop retained variant | 71 | 58 | 28 | 15 |
| coding sequence variant | 13 | 11 | 6 | 4 |
| 5 prime UTR variant | 5966 | 5970 | 2089 | 2093 |
| 3 prime UTR variant | 10976 | 11165 | 3007 | 3196 |
| non coding transcript exon variant | 2301 | 2420 | 823 | 942 |
| intron variant | 9808 | 9630 | 2619 | 2441 |
| non coding transcript variant | 2375 | 2495 | 826 | 946 |
| upstream gene variant | 41520 | 41569 | 9515 | 9564 |
| downstream gene variant | 43158 | 43301 | 9853 | 9996 |
| intergenic variant | 1 | 1 | 0 | 0 |

We found 4,479 Pokkali and 4,518 IR29 genes with overlapping SNPs that were also differentially expressed under salt stress. Both varieties contained SNPs in the *SalT*, *DREB*, *RING,* and *NAC* genes. SNPs were also found in members of formin-like protein-coding genes. Formin-like proteins are involved in rice growth and development (Yang et al., 2011; Zhang et al., 2011). SNPs unique to Pokkali were also identified in the differentially expressed gene *RBOHB*. SNPs were identified in embryonic abundant protein-coding genes in IR29. These SNPs may have potential consequences for the ability of IR29 to protect against protein aggregation during salt stress [49]. Eight genes with unknown functions carrying the polymorphic SNP colocalize the Saltol QTL region. InterProscan (Hunter et al., 2012) predicted half of these genes to contain DnaJ domains or domains of unknown function (DUF). Among the genes overlapping the 257 polymorphic SNP sites, 42 and 37 genes were differentially expressed in Pokkali and IR29, respectively (Supplementary Table 8).

## Discussion

Rice is cultivated in river basins, uplands, lowlands, and near coastal areas. However, due to extensive farming, native agronomic practices, and weather-related events such as hurricanes in the coastal areas, the growth environment, including the soil, table water, and flooding, often leads to hypersaline conditions. To find reliable candidate genes with some assessment of their function, expression, and associated genetic markers, we de novo assembled and characterized the transcriptomes of two rice varieties in response to salt stress. We used two O. sativa indica lines for comparison: the salt-sensitive IR29 breeding line and a salt-tolerant Pokkali landrace. Salt-tolerant landraces are still commonly used by farmers; however, these landraces also contain traits such as low yield that may be counterproductive for cultivation. Advancements in molecular breeding methods aim to understand the genetic control of salt tolerance mechanisms. This transcriptome-based study is a rice resource that (1) provides information on candidate genes and markers for improving salt tolerance traits, (2) provides data that extends existing resources to study in-depth molecular events and their perturbations under salt stress, and (3) helps improve reference genome annotations by identifying novel salt-responsive alternatively spliced transcript isoforms and transcribed regions of the two genomes.

### Gene expression within the Saltol QTL

Various genetic studies on rice have identified salt sensitivity QTLs span the genomic regions of chromosomes 1, 3, 4, 5, 6, 7, and 9. Of these, the major salt tolerance QTL *Saltol* mapped on chromosome 1 with genes involved in regulating the sodium-potassium ratio and salt-tolerance breeding [74–77]. In our study, we identified 85 differentially expressed genes located within the Saltol QTL [77]. Most notable among the set are four genes, *SalT*, *OsBZ*8, *Respiratory Burst Oxidase Homolog B* (*RBOHB*, OS01G0360200; Pokkali_10699, IR29_03274, OS01T0360200-01), and an *Ethylene response factor* (ERF) (Figure 5 and Supplementary Figure S2). The *SalT* gene (OS01G0348900) was upregulated in the salt-tolerant Pokkali at 2 and 5 hours after exposure to salt, with transcript abundance staying above those of untreated samples at the same time points. However, in IR29, the *SalT* gene downregulated at 5 and 10 hours post-salt exposure, with abundance staying much lower than that in the untreated samples. The *SalT* gene is known to interact with *OsBZ8* [78], a salt stress-related transcription factor discussed earlier. In Pokkali, we found that *RBOHB* remained upregulated at 1, 2, and 5 hours. Respiratory burst oxidases have been shown to generate extracellular superoxide in response to pathogens [79] and may play a more general role in stress response. In response to oxidative burst generation, plants utilize glutaredoxins to avoid biological damage [80]. We found that *glutaredoxin-C1* (*GRXC1,* Os01G0368900) was downregulated in IR29 at 2 hours and Pokkali at 10 hours. This downregulated expression pattern of glutaredoxins in rice is similar in response to cold stress [5]. Another notable gene, OS01G0367900, overlapping the Saltol QTL, encodes a *Chromatin-remodeling complex ATPase chain*. This gene was only upregulated at 5 hours in Pokkali, but its overall expression stayed much lower than that of IR29, suggesting that it may be involved in regulating gene expression [81].

**Figure 5.**
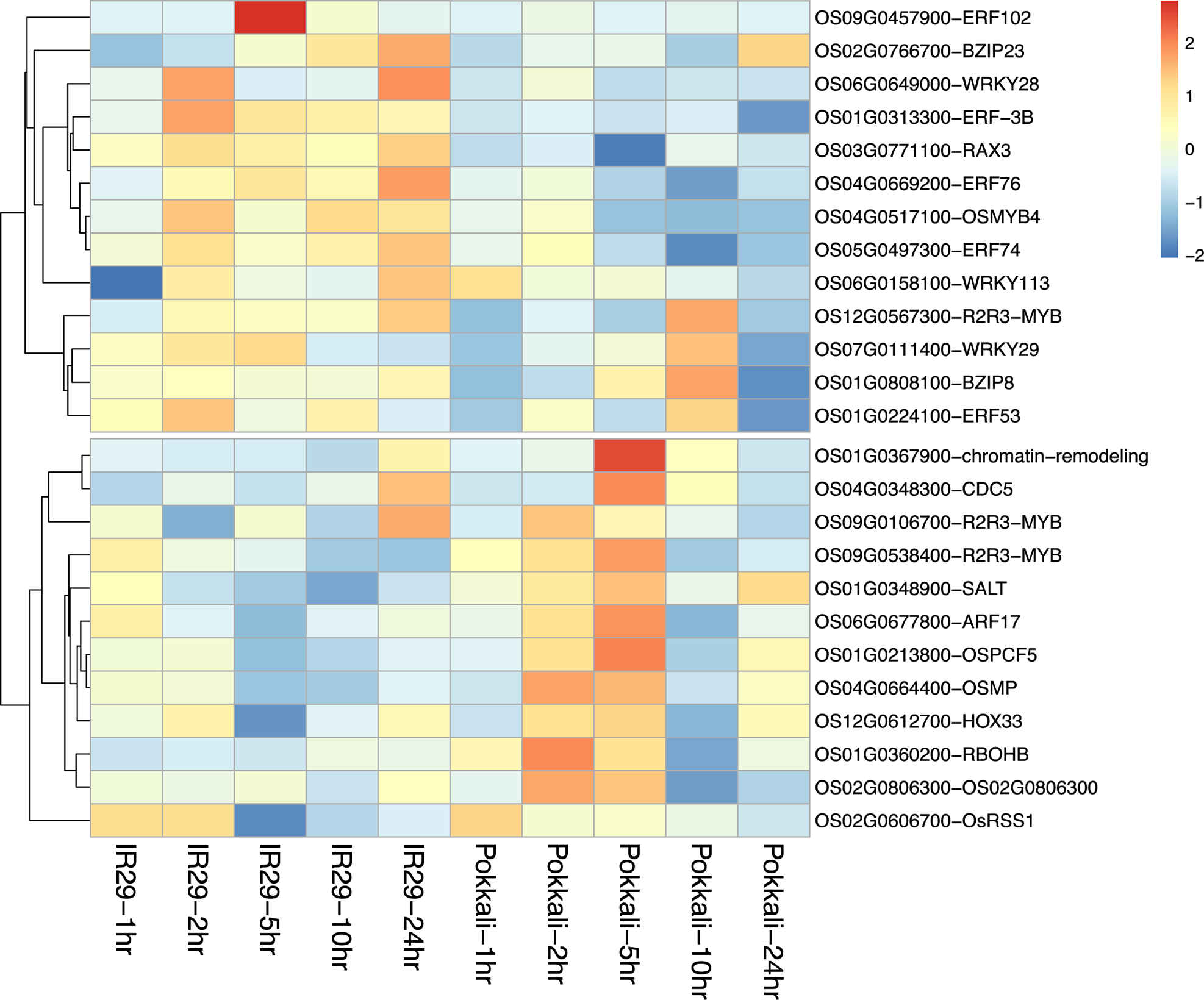
Heatmap of coregulated transcription factors targeting salt-stress responsive genes. Rows represent transcription factors targeting the gene, and columns represent salt experiment time points (1, 2, 5, 10, and 24 hours). Individual cells represent the fold-change expression values for the transcription factor at a specific time point.

### Identification of putative transcription factor-binding sites upstream of salt-stress responsive genes and coregulated transcription factor gene targets

Only Pokkali contained differentially expressed transcription factors within the Saltol QTL. *Ethylene response factor 3B* (*ERF-3B*, OS01G0313300) transcription factor was downregulated in response to salt stress at all five-time points in Pokkali (Figure 5). ERFs are known to contribute to tolerance to salt stress by regulating the expression of oxidative stress genes [82–84].

We also identified transcription factors (TFs) that have putative binding sites in the 1 kb upstream region of protein-coding genes. Currently, there is limited knowledge of PWMs for rice transcription factors. Since Arabidopsis has a comprehensive list of PWMs for many transcription factor families available in the JASPAR and TRANSFAC databases, we used the Arabidopsis-rice orthology datasets [85, 86] to select PWMs appropriate for use in rice. Using data generated from this TFBS analysis and by filtering the list of genes that are differentially expressed in the Pokkali and IR29 rice varieties, we computationally analyzed potential regulatory relationships between rice transcription factors and their target genes, including *SalT*, *OsBZ8*, *RBOHB,* chromatin remodeling genes from the *Saltol* QTL region on Chromosme-1 and *RSS1* on Chromosome-2. We also analyzed coexpression relationships between transcription factors and their target genes (Figure 5). These genes group into two distinct clusters. Genes in cluster #1 were upregulated at 2-24 hour time points in IR29 versus the cluster #2 genes showing preferred upregulation at 2 and 5 hours in Pokkali. The cluster #1 genes include various ethylene response factors (ERFs), *WRKY,* and the target *BZIP8*. The cluster #2 genes *HOX3, PCF5, R2R3-MYB CDC5, ARF15,* and *OSMP* transcription factors were upregulated at 2 and 5 hours along with their targets *SALT*, *RBOHB,* and *chromatin remodeling* genes. The cluster #2 genes in Pokkali were either expressed in low amounts at 1 and 24 hr or downregulated overall in salt-susceptible IR29. It suggests that these genes in Pokkali and IR29 respond oppositely under salinity stress.

### Identification of SSR markers overlapping the *RBOHB* and *RSS1* genes

To look for genetic markers (SSRs and SNPs) with consequences for differential expression, gene function, and gene structure, we identified the presence of polymorphic SSRs in the salt-responsive genes *RBOHB* and *RSS1*. These polymorphic SSRs were present in the 5’-UTR of *RBOHB* (Supplementary Figure S8) and the 3’-UTR of *RSS1* (Supplementary Figure S9). 5’-UTRs carry ribosome binding sites, function in transcript stability, provide translational efficiency, and may affect transcript abundance. Similarly, 3’-UTRs have a role in providing transcript stability that may affect transcript abundance. Previous studies suggest SSRs may contribute to advantageous mutations and be associated with phenotypic traits in plants [87] and humans [88, 89].

The *RBOHB* 5’-UTR SSR in Pokkali has (GA)9 dinucleotide repeats compared to longer (GA)13 repeats in IR29. We examined the predicted RNA fold structure of the SSR containing *RBOHB* transcripts from Pokkali and IR29. Without six specific nucleotides, Pokkali loses an internal loop, whereas IR29 forms a longer stem and an additional internal loop (Figure 6). We hypothesize that the length of the IR29 SSR at the 5’ end leads to transcript instability and faster turnover (yet to be tested), potentially appearing as a downregulation of *RBOHB* (measured by transcript abundance) in our expression data. The *RBOHB* SSR in our data is the same as the rice SSR genetic marker RM10887 [90].

**Figure 6.**
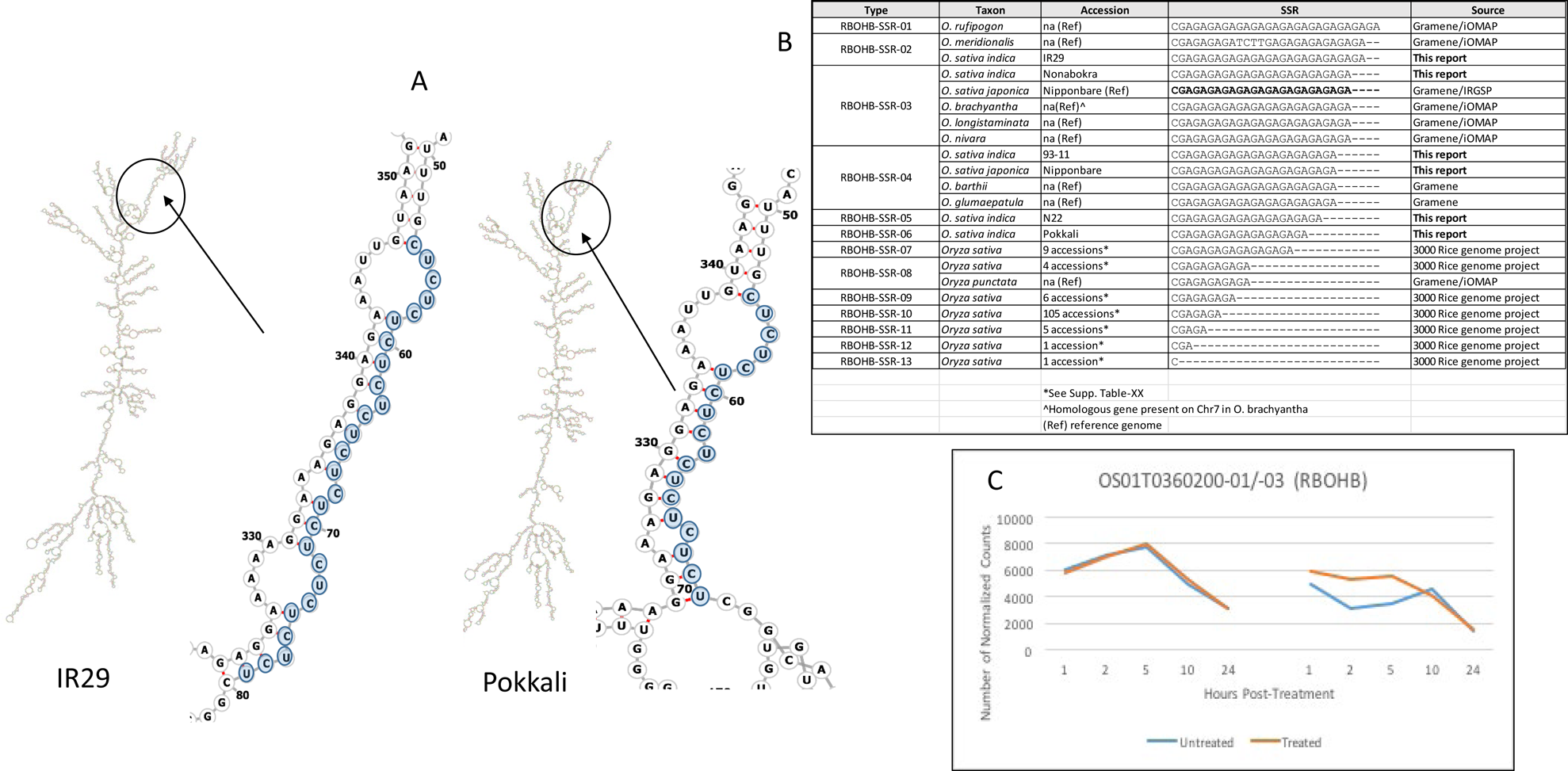
*RBOHB* mRNA secondary structures. Enhanced images show the position of the SSR in the structure (A). Alignments of the repeat region sequences in multiple rice lines and various species of *Oryza* (B). Line graphs showing normalized read counts (transcript abundance) for the *RBOHB* gene in respective rice lines at different time points.

Similarly, analysis of the trinucleotide (GTA)n repeat SSR region found in the *RSS1* 3’-UTR suggests the formation of a shorter hairpin loop structure in Pokkali RNA (Figure 7). A shorter hairpin structure is known for thermodynamic stability, which may contribute to better transcript stability that may be the reason for higher mRNA abundance in Pokkali during the 2-5 hour time period compared to IR29, which showed a rapid change in transcript abundance [91, 92]. These observations provide novel insights, and additional studies are needed to validate the hypothesis.

**Figure 7.**
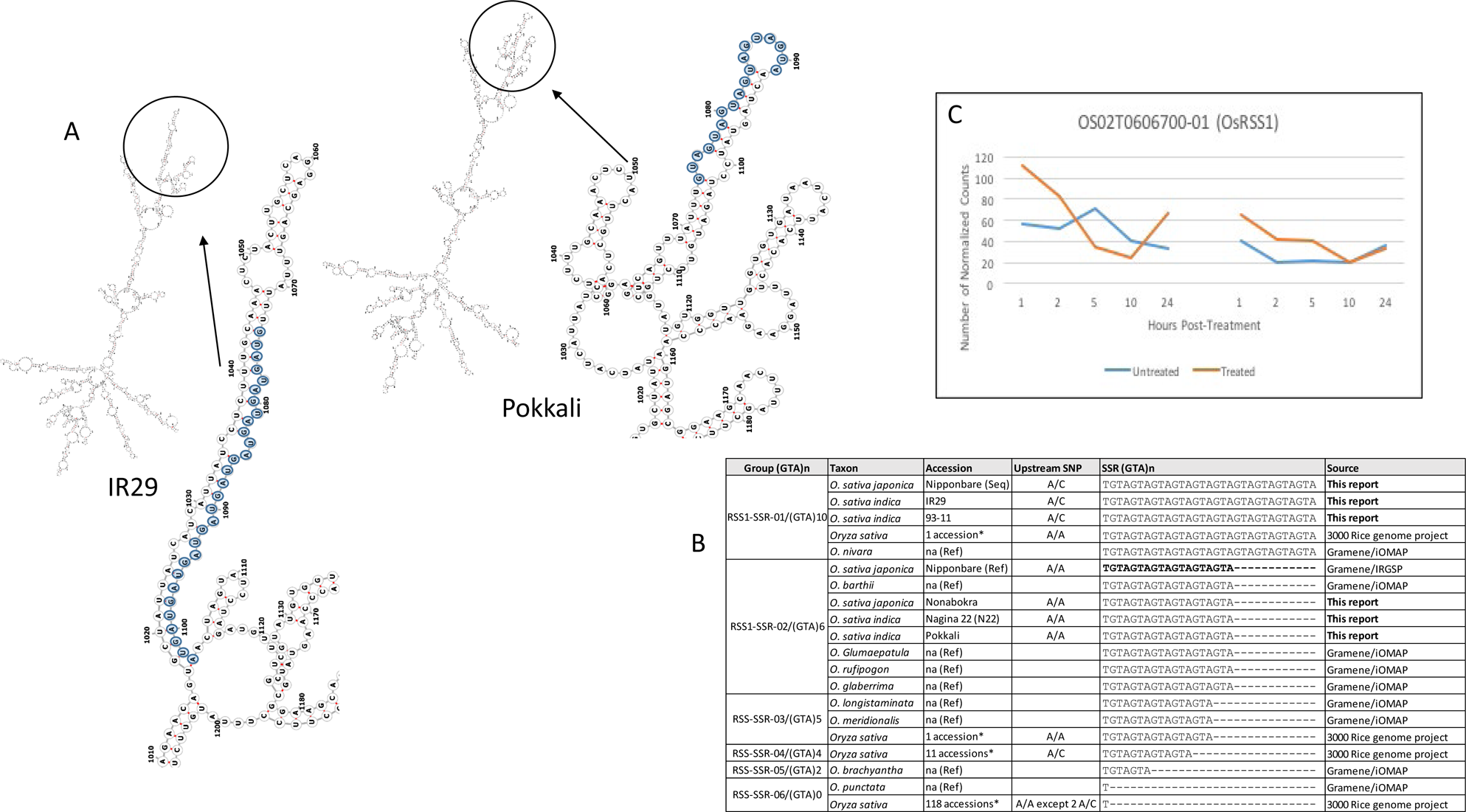
*RSS1* mRNA secondary structures. Enhanced images show the position of the SSR in the structure (A). Alignments of the repeat region sequences in multiple rice lines and various species of *Oryza* (B). Line graphs showing normalized read counts (transcript abundance) for the *RSS1* gene in respective rice lines at different time points.

To confirm the presence of polymorphic SSRs in *RBOHB* and *RSS1* UTR sites and explore the genetic diversity, we sequenced the PCR-amplified fragment of the respective genome region from a number of indica and japonica rice lines. We also analyzed the genetic diversity data from the 3000 Rice Genome Project [93]. These sequences were aligned against the reference genomes of japonica (Nipponbare), indica 93-11, and wild *Oryza* species. In our panel, Nagina 22 (N22) is a drought- and heat-tolerant AUS variety and Nona Bokra is a salt-tolerant indica variety. For *RBOHB,* we observed 13 different variety clusters based on their SSR length (Figure 6). N22 has the next shortest CU repeat compared to Pokkali, whereas Nona Bokra contains a CU repeat almost as long as IR29. For *RSS1*, N22 and Nona Bokra shared the GUA repeat with the same length as in Pokkali (Figure 7). While this result was expected between Pokkali and Nona Bokra with *RSS1*, our results suggest that N22 may also be a candidate for salt tolerance studies.

Our Sanger sequence validation data of *RSS1* also revealed a C to A SNP 27 base pairs upstream of the targeted repeat in the IR29 and 93-11 lines. We also identified this particular SNP in our variant effect production analysis which called it a 3’ UTR variant. Further investigation of this variant marker may reveal a possible association with salt-tolerant traits.

### Improving genetic resources

Using publicly available bioinformatics tools, databases, and openly accessible FAIR datasets enabled us to analyze high-quality transcriptomes of the IR29 and Pokkali rice varieties in response to salt stress. We chose to make our comparisons to the *O. sativa* japonica group due to its abundance of annotation resources. Approximately 3% of our transcriptome assemblies did not map to currently annotated gene models but did map to the reference genome. Some of our assembled transcripts revealed structural differences compared to the reference gene models (e.g., Supplementary Figures S8 and S9). It suggests that these transcripts may be novel isoform variants that are yet to be annotated in the reference Japonica genome. Future studies will explore the differences between de novo assemblies and reference gene models.

This study provides genomic resources that include transcriptome assemblies, their functional annotation, differentially expressed genes, and the identification of potential genetic markers from the two rice varieties. The number of time points sampled in this analysis makes these data valuable in providing a detailed timeline of how salt-tolerant and salt-sensitive rice varieties respond to salt stress. This study further provides a resource for identifying novel genes, transcript models, and coexpression networks that can ultimately yield insight into improving a crop essential to feeding the world population.

## Authors and contributors

PJ conceived and led the project. SF and AS carried out the salt treatment experiment. SF and AS extracted RNA. MG and SF conducted de novo transcriptome assembly, various blast analyses, annotation, differential gene expression analysis, transcription factor mining, SSR analysis, SNP identification, and variant consequence analysis. MA and MM conducted transcription factor-binding site identification. MG, SF, and PJ wrote the manuscript. All authors have read, edited, and approved the manuscript.

## Acknowledgments

The authors would like to thank Dr. Mamatha Hanumappa for help in setting up the experiment, the Center for Genome Research and Biocomputing core facility staff Anne-Marie Girard and Caprice Rosato for the quantity and quality assessment of RNA, Mark Dasenko for Illumina sequencing, and Christopher Sullivan for computational support. We also appreciate the complimentary access to CropPedia provided by Anker Sørensen from Keygene to explore the 3000 rice genome project data.

## Supplementary material

**Supplementary Figure S1.**

A. Multidimensional scaling (MDS) analysis of RNA-Seq reads for IR29 samples.

B. Multidimensional scaling (MDS) analysis of RNA-Seq reads for Pokkali samples.

C. Multidimensional scaling (MDS) analysis of RNA-Seq reads for all samples.

**Supplementary Figure S2a and S2b.**

Line graphs displaying the number of normalized read counts of featured genes described in the manuscript.

**Supplementary Figure S3.**

A. Analyses of differentially expressed transcripts over 5 time points.

B. The four-way Venn diagrams show the distribution and commonalities of Japonica homolog counts with respect to their response postsalt exposure in IR29 and Pokkali. The bar graph shows the number of unique transcripts that are differentially expressed at each time point in IR29 and Pokkali.

**Supplementary Figure S4.**

The frequency distribution of transcripts of varying sizes in the de novo transcriptomes of IR29 and Pokkali and their comparison with annotated transcripts of japonica rice reference genome.

**Supplementary Figure S5.**

Gene Ontology IDs associated with the de novo transcriptomes of IR29 and Pokkali.

**Supplementary Figure S6.**

Number of SSRs identified in the de novo transcriptomes of IR29 and Pokkali.

**Supplementary Figure S7.**

SNP counts by chromosome

**Supplementary Figure S8.**

View of RBOHB SSR and the aligned transcripts on the Gramene genome browser.

**Supplementary Figure S9.**

View of the RSS1 SSR and the aligned transcripts on the Gramene genome browser.

**Supplementary Table 1.**

Normalized counts of differentially expressed genes in IR29 and Pokkali. Data includes counts in each replicate of control and salt treated samples at each timepoint.

**Supplementary Table 2.**

List of differentially expressed Japonica genes that encode transcription factors. Data includes gene expression patterns (red for upregulation, blue for downregulation and white for N/A)

**Supplementary Table 3**

Unique GO terms identified in IR29 and Pokkali transcript annotations

**Supplementary Table 4.**

List of differentially expressed Japonica genes near assembled transcripts aligned to the Japonica genome but not gene models.

**Supplementary Table 5.**

Counts of simple sequence repeats identified in IR29 and Pokkali transcriptome data.

**Supplementary Table 6.**

Position of the 257 polymorphic SNP set on the Japonica genome.

**Supplementary Table 7.**

Number of transitions and transversions resulting from SNP analysis reference to Japonica genome.

**Supplementary Table 8.**

List of Japonica genes overlapping the 257 polymorphic SNP set

