## Supplementary material for "Comparative Analysis of Salinity Response Transcriptomes in Salt-Tolerant Pokkali and Susceptible IR29 Rice": Geniza_etal_supplementary_figures.pdf

A B

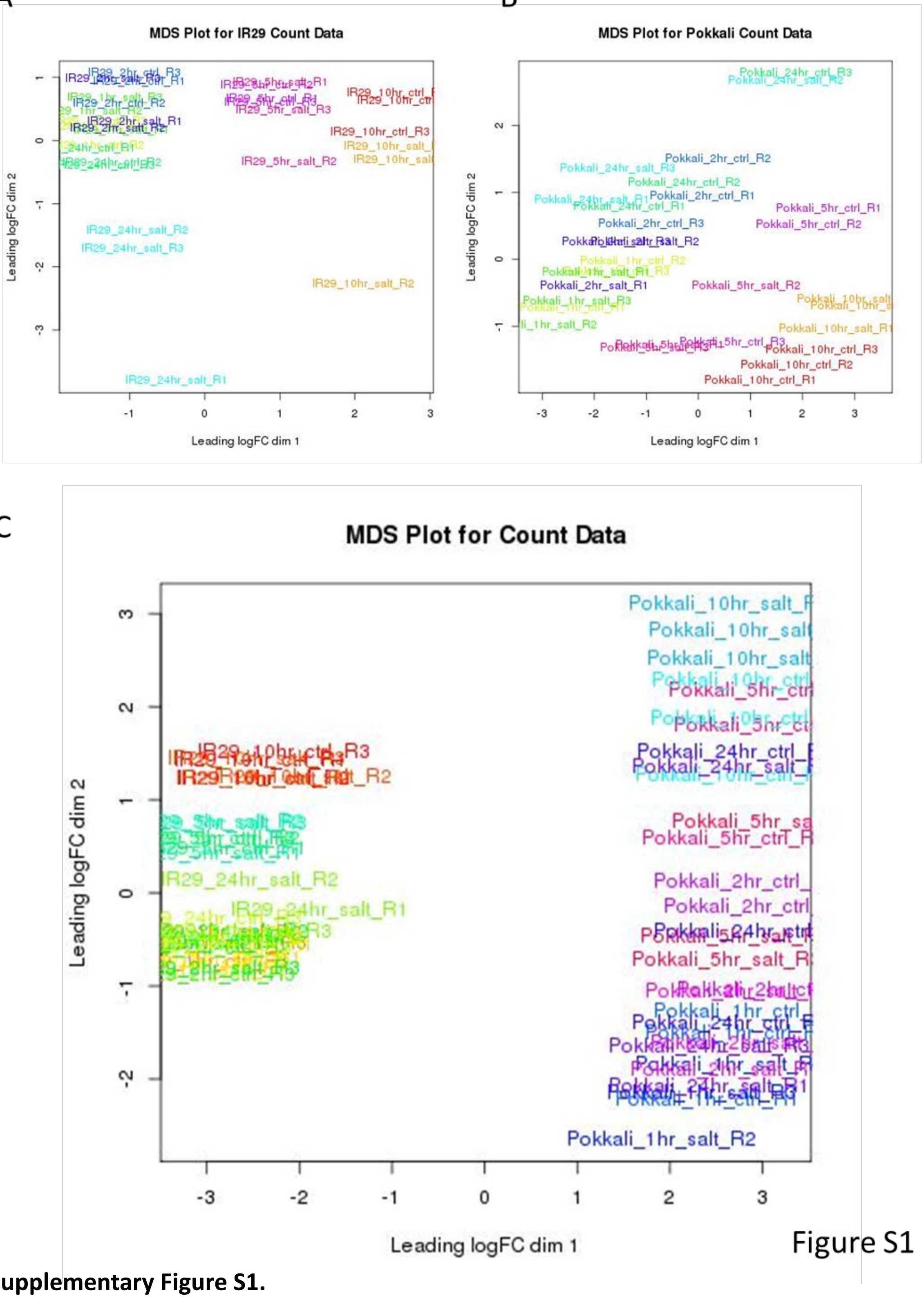


C

Figure S1

**Supplementary Figure S1.**

A. Multidimensional scaling (MDS) analysis of RNA-Seq reads for IR29 samples.

B. Multidimensional scaling (MDS) analysis of RNA-Seq reads for Pokkali samples.

C. Multidimensional scaling (MDS) analysis of RNA-Seq reads for all samples.

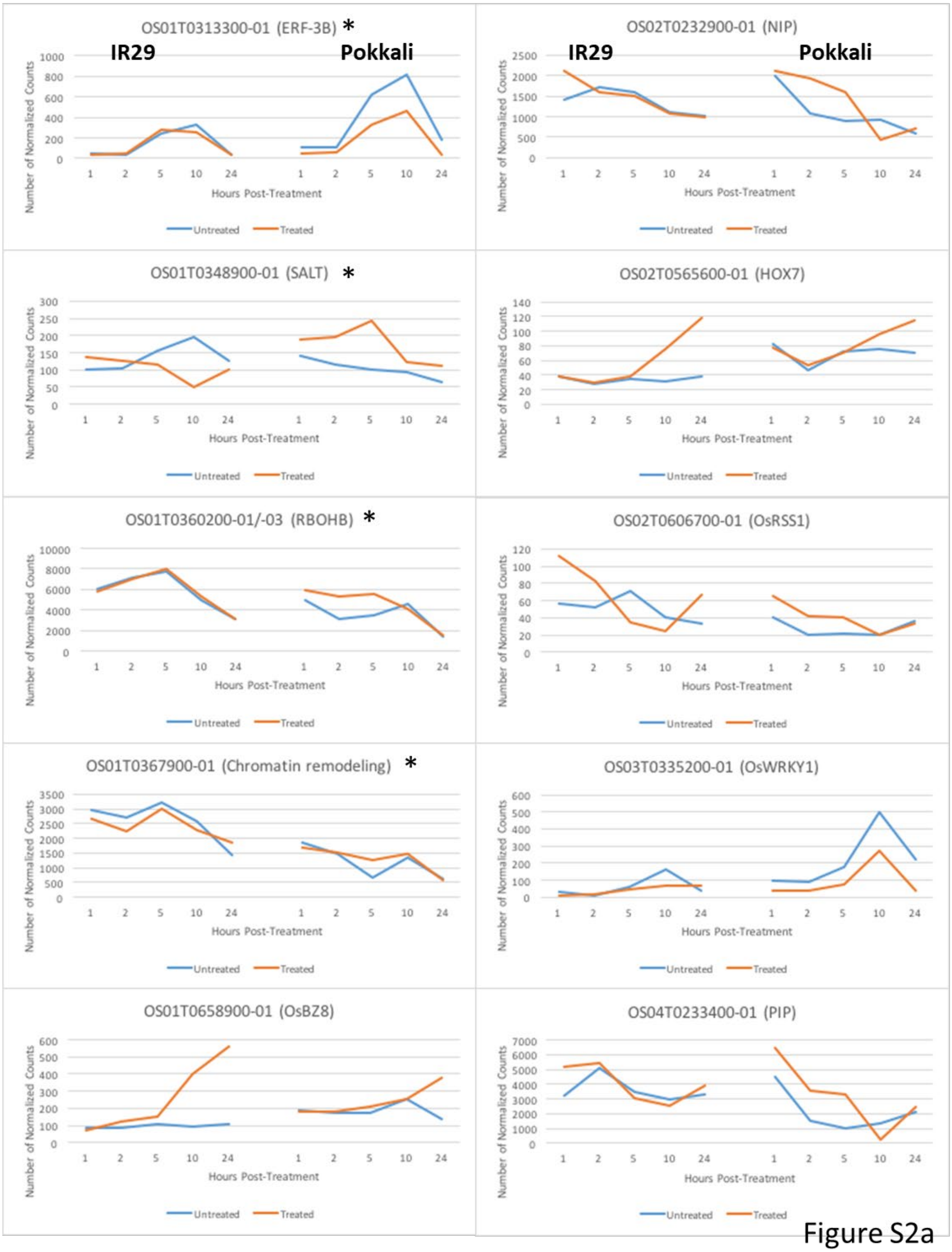


Figure S2a

**Supplementary Figure 2a and 2b.**

Line graphs displaying the relative count number of featured genes in the manuscript.

* = within the Saltol qtl

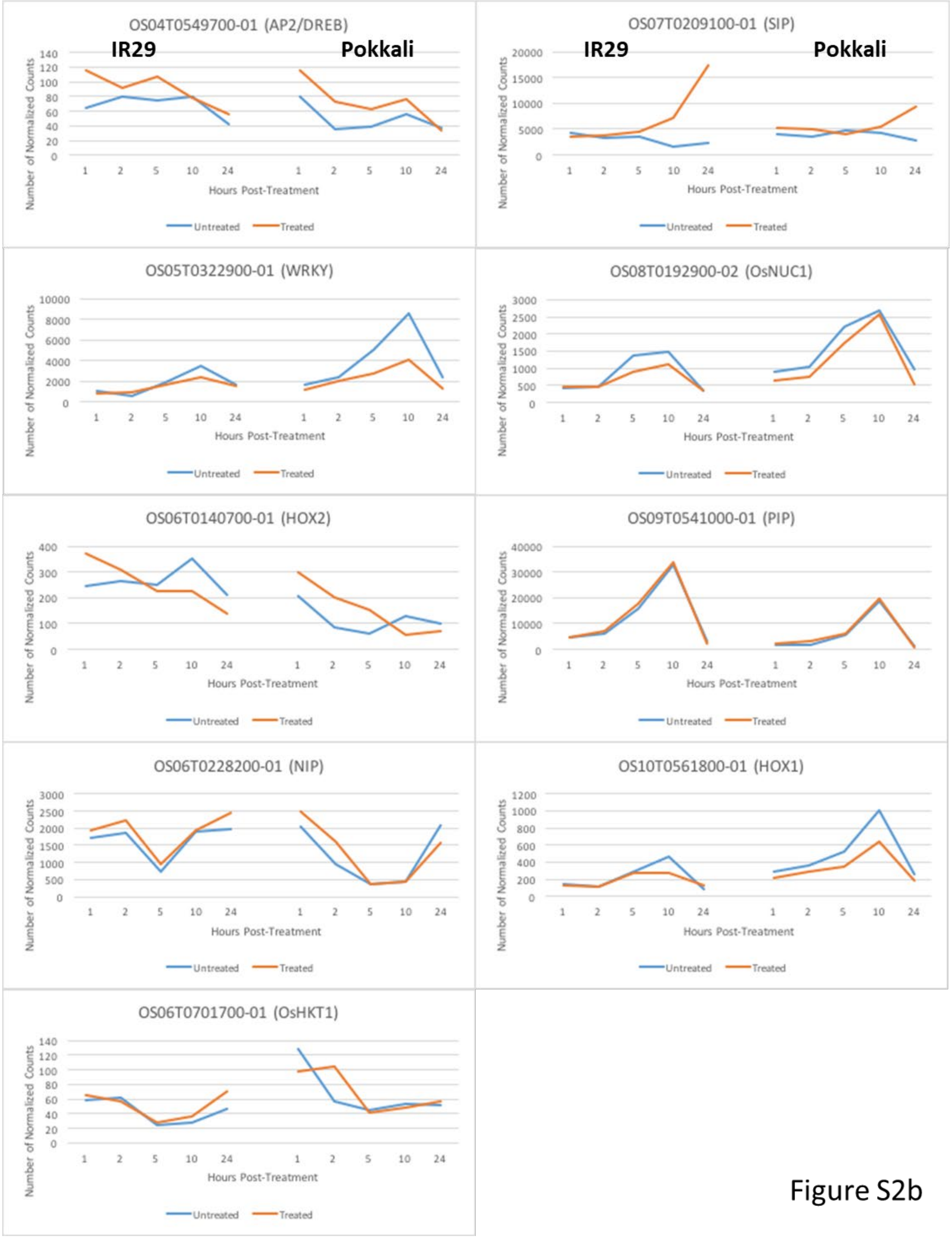


Figure S2b

**Supplementary Figure 3a and 3b.**
Line graphs displaying the relative count number of featured genes in the manuscript.
* = within the Saltol qtl

Figure S3

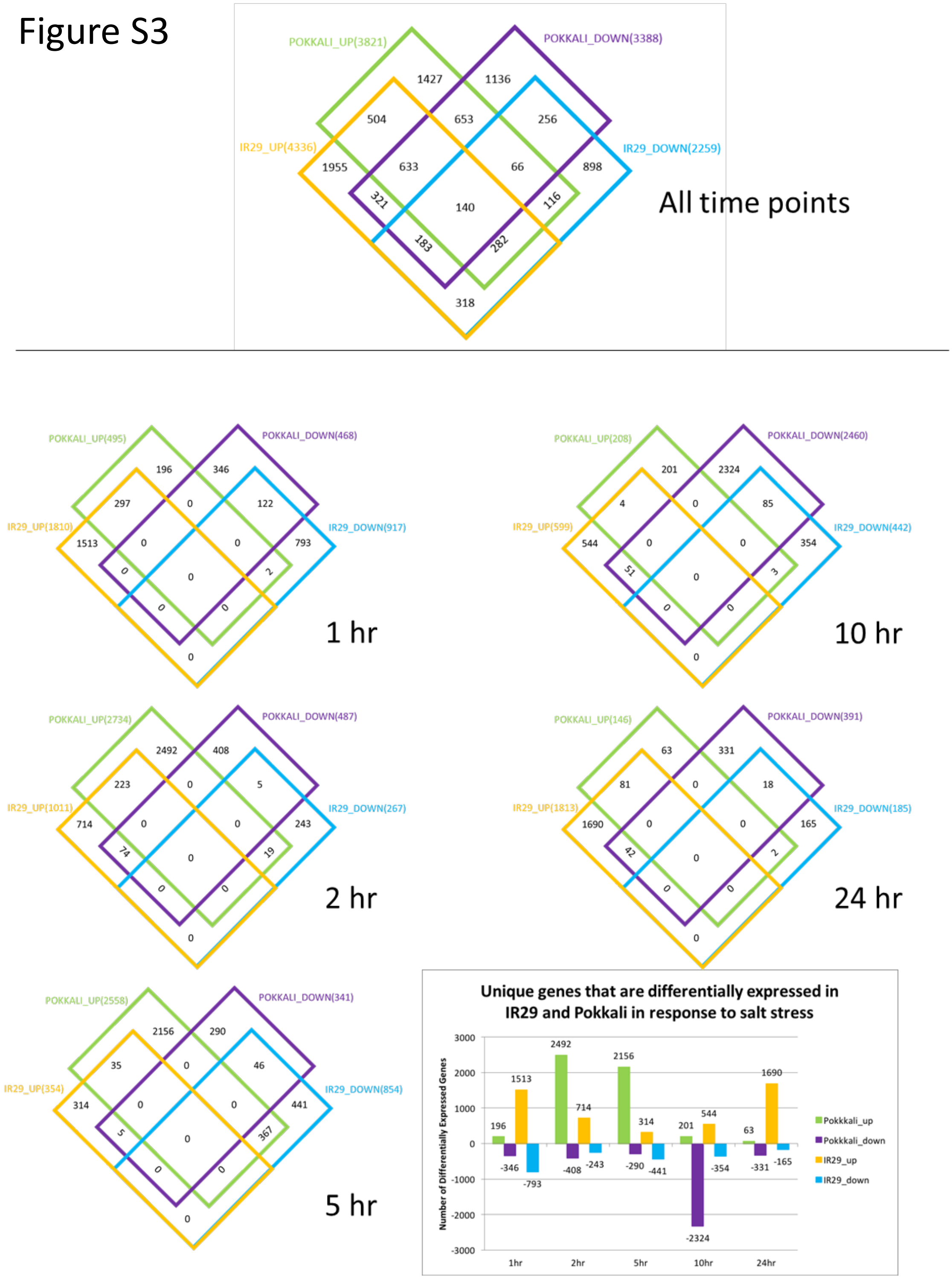


**Supplementary Figure S3.**

A. Analyses of differentially expressed transcripts over 5 time points.

B. The four-way Venn diagrams show the distribution and commonalities of Japonica homolog counts with respect to their response postsalt exposure in IR29 and Pokkali. The bar graph shows the number of unique transcripts that are differentially expressed at each time point in IR29 and Pokkali.

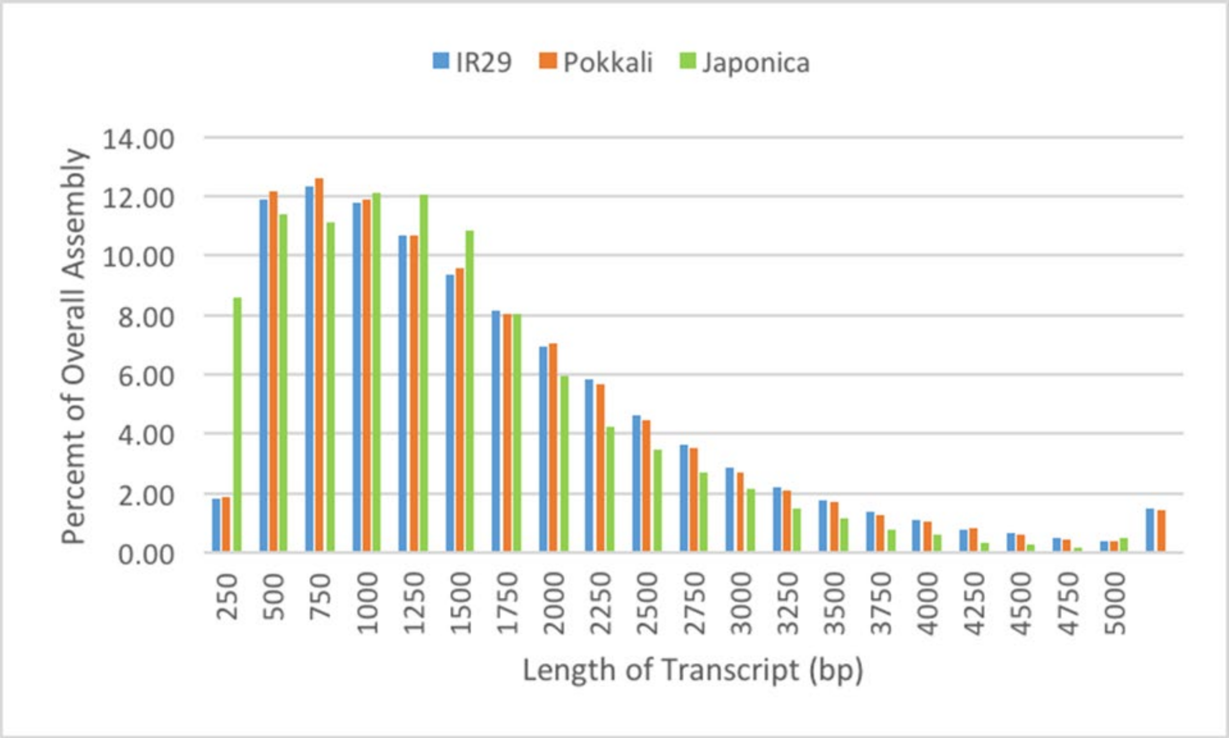


Figure S4

**Supplementary Figure S4.**
The frequency distribution of transcripts of varying size in the de novo transcriptomes of IR29, Pokkali, and the annotated transcriptomes of Japonica.

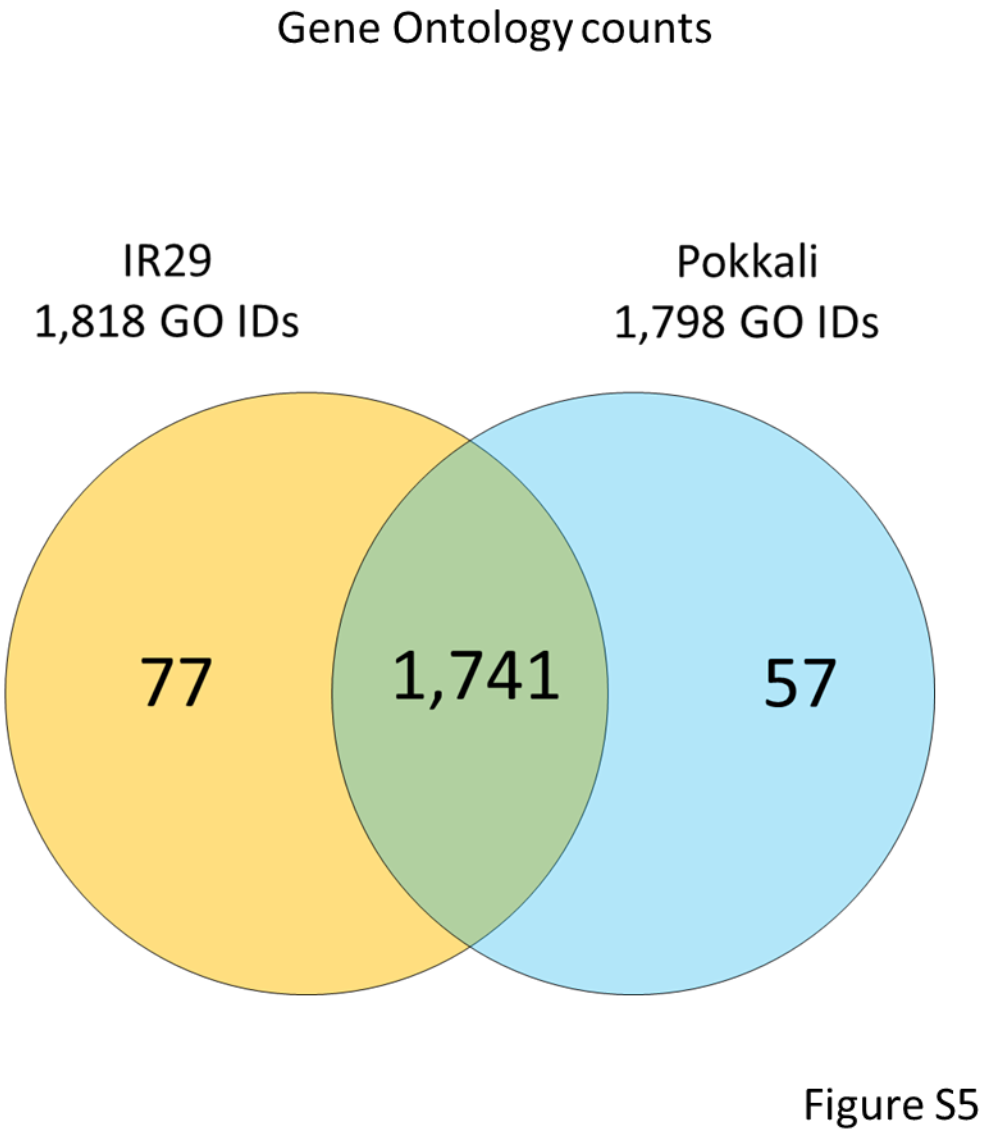


**Supplementary Figure S5.**
Gene Ontology IDs associated with the de novo transcriptomes of IR29 and Pokkali.

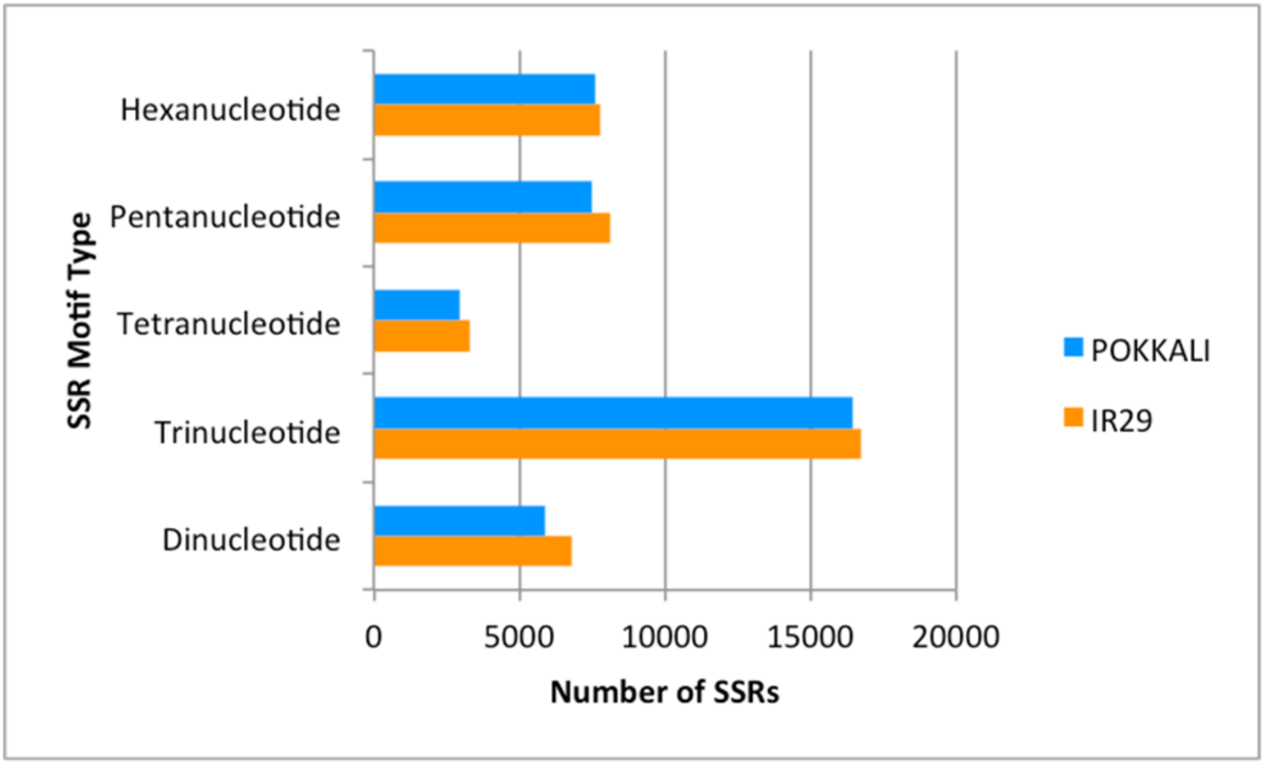


Figure S6

**Supplementary Figure S6.**

Number of SSRs identified in the de novo transcriptomes of IR29 and Pokkali.

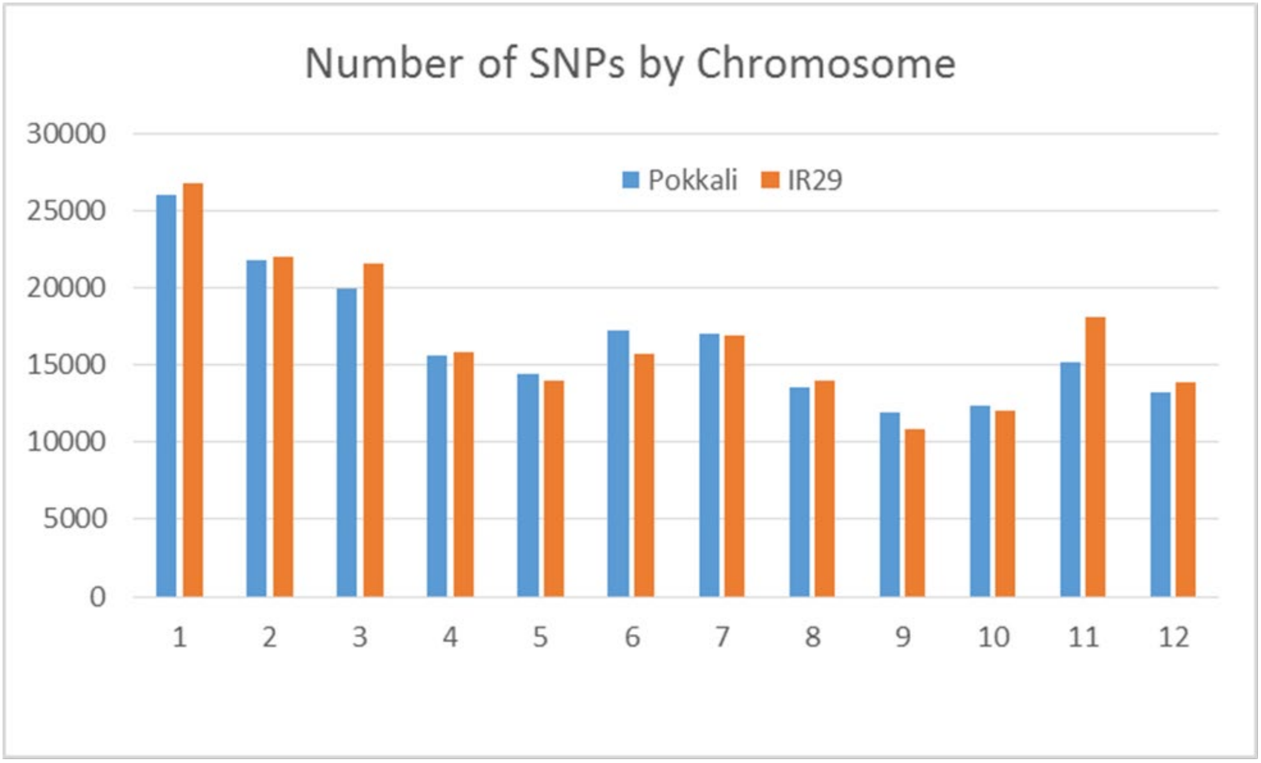


Figure S7

**Supplementary Figure S7.**

SNP counts by chromosome

### RBOHB / OS01G0360200

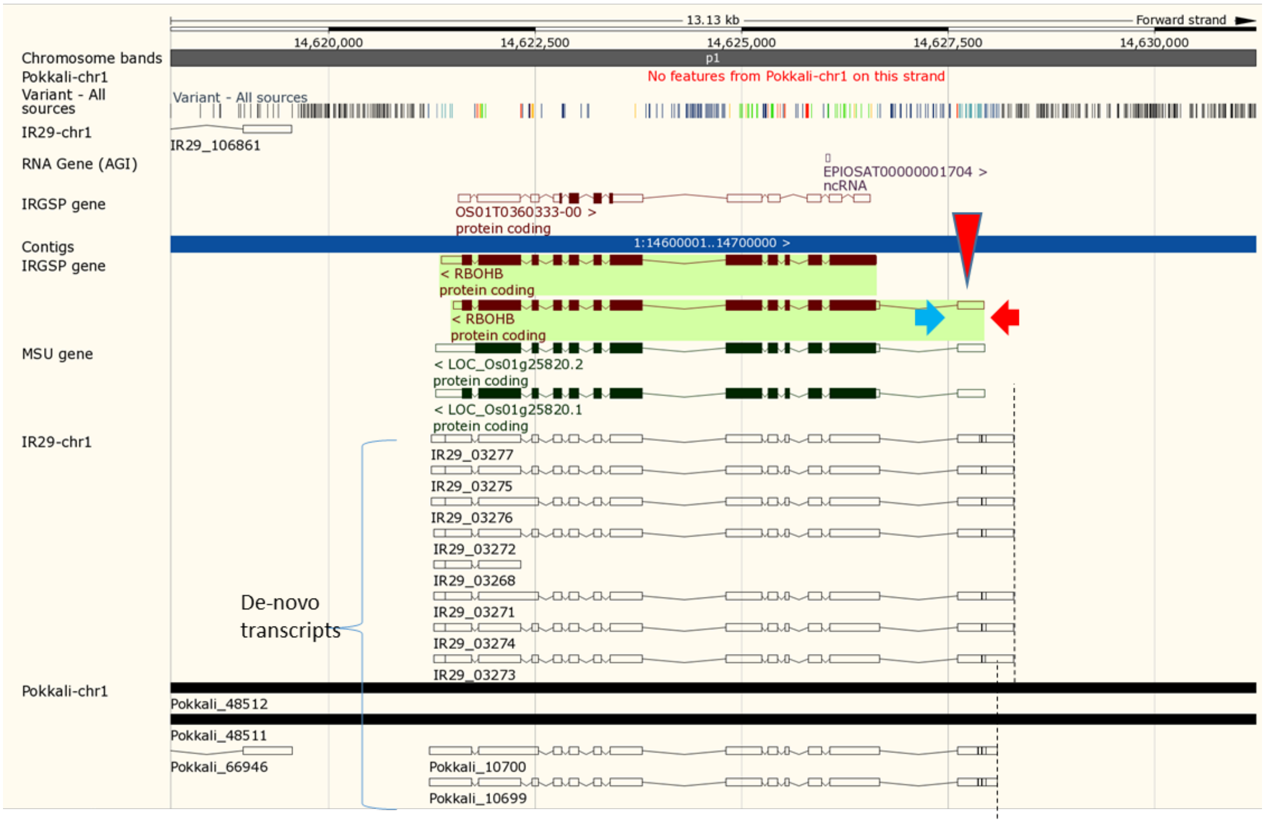


RBOHB-FWD:

5' – GCG ACC ACA AAA AGC TGG AG – 3'

RBOHB-REV:

5' – TCA AGC TGT CCA AAC ACC GT – 3'

SSR

Forward primer

Reverse primer

Figure S8

**Supplementary Figure S8.**
View of RBOHB SSR and the aligned transcripts on Gramene genome browser.

### RSS1 / OS02G0606700

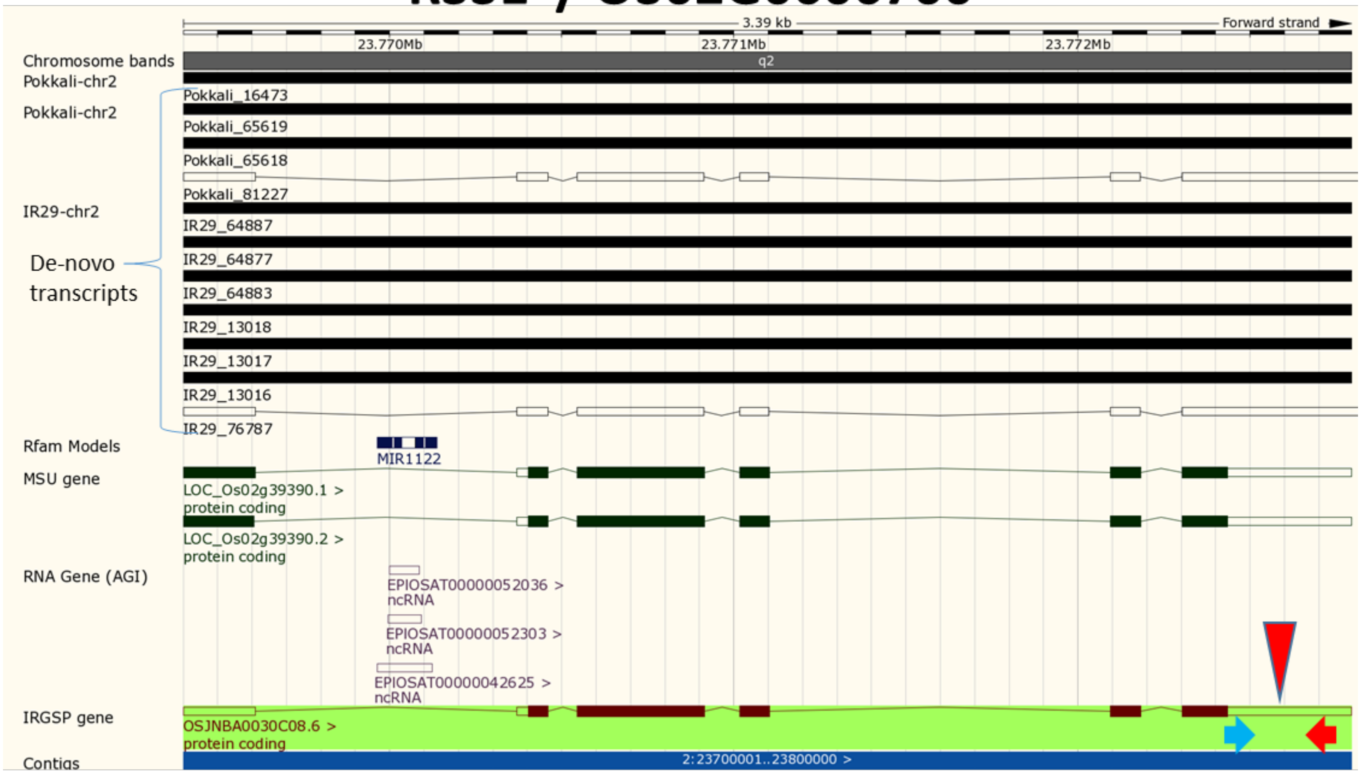


RSS1-FWD:
5’ – ACT CCT GGA GCC TGG AAT GA – 3’
RSS1-REV:
5’ – TTG CTT CCG CTA CTT GGG TT – 3’

SSR
Forward primer
Reverse primer

Figure S9

**Supplementary Figure S9.**
View of the RSS1 SSR and the aligned transcripts on the Gramene genome browser.
